# FLASH-P: Turning decades of biology into accurate causal networks with AI agents

**DOI:** 10.64898/2026.06.13.731799

**Authors:** Christos Mitsanis, Nicole Fortuna, Christine Beveridge, David Kainer

**Affiliations:** School of Agriculture and Food Sustainability Faculty of Science, The University of Queensland, Queensland, Australia; Queensland Alliance for Agriculture and Food Innovation, The University of Queensland, Queensland, Australia; Australian Research Council Centre of Excellence for Plant Success in Nature and Agriculture, The University of Queensland, Queensland, Australia

## Abstract

Mechanistic networks that encode causal regulatory logic can predict the effects of genetic and environmental perturbations but constructing them is a bottleneck in systems biology because the relevant knowledge lies scattered across thousands of resources, untapped for both building and validating such networks. Here we present FLASH-P, a multi-agent framework that autonomously curates this literature into perturbable, signed-directed network models for any trait–species combination in under an hour without much computational power. Twelve FLASH-P networks across seven species predicted the directional outcome of 1,088 published perturbations with a mean accuracy of 90%. This accuracy was driven by the regulatory topology FLASH-P constructs, which is why it outperformed knowledge-graph derived networks. Its merging agent combined six networks into one that preserved single-trait accuracy and recovered pleiotropic effects, and consolidated independent runs of one trait into a comprehensive, high-accuracy network. FLASH-P networks enable applications that require trait models.

## Main

Phenotype emerges from the complex interplay among molecular entities (genes, RNAs, proteins, metabolites) and external environmental influences. These interactions form the basis of mechanistic networks that model the influences and effects on a given trait. Mechanistic networks encode biological causal logic that can be used to predict the dynamic consequences of perturbations such as gene knockouts, exogenous supply of compounds, and environmental changes. At scale and with sufficient accuracy, such networks would constitute powerful tools across biomedicine, bioengineering and possibly agriculture, as already demonstrated by quantitative systems pharmacology in drug development^1^ and by genome-scale metabolic models in industrial yeast strain design^2,3^. Such network construction is slow, and the methodology is unsettled, so producing an accurate mechanistic network even for a single well-researched trait represents one of the great challenges in systems biology^4^.

One approach is to experimentally generate data (often single-cell gene expression), both with and without perturbations, while measuring particular phenotypes and environmental variables. A network is then constructed and parameterized from the observations using software such as RENGE^5^, scDNS^6^, TxPert^7^, MAVEN^8^, PDGrapher^9^. However, perturbation screens using technologies like perturb-seq or CRISPR multiplexing are invariably resource heavy, and in plants they face efficiency and scaling problems^10,11^. Stable transformation takes weeks to months, accurate phenotyping usually requires growing plants to maturity, and hence most current perturbation software is heavily biased toward mammalian data. Consequently, few mechanistic networks are constructed in plants, despite a clear potential role in prioritising candidate genes for crop breeding, predicting the effects of gene stacking on whole-plant phenotype, transferring knowledge to less-studied species, and accelerating hypothesis generation in fundamental plant science.

The gap matters most for genomic prediction, where statistical models^12^ learn statistical correlations between markers and traits rather than the underlying biology. A mechanistic network supplies that missing causal layer, enabling prediction for combinations of genotypes and environments not seen in training, which is the central aim of the Breeding 4.0 paradigm ^13^.

Yet mass perturbation data in plants already exists. Decades of plant molecular experimentation has produced thousands of individual gene knockout, overexpression, and hormone application studies across a multitude of species. Each study typically characterises a small number of perturbations in depth and reports the phenotypic and molecular consequences. Collectively, this body of work represents an enormous perturbation matrix fragmented across thousands of papers, albeit with inconsistent reporting formats, varying genetic backgrounds, varying environments and experimental designs and diverse types of recorded observation. No single experimental campaign could viably reproduce this breadth of interventions, traits and species, so how can we make effective use of all this prior information?

Here we introduce a multi-agent framework for autonomously generating high-quality, perturbable mechanistic networks of a target trait and species from prior knowledge alone. Our approach, FLASH-P, builds upon recent advances in the reasoning capabilities of frontier large language models (LLMs), coupled with agentic controls and constraints. FLASH-P exploits the fact that for many traits, precise regulatory information exists piecemeal in the literature and could be engineered into a highly accurate mechanistic network if curated and synthesised intelligently. When a user prompts FLASH-P with a trait-species combination (*e.g.* “kernel number in Maize”), a team of autonomous AI agents researches and curates a causal network that links genes, transcription factors, metabolites, processes, and environmental influences on each other and, via causal chains, to the target trait. The agentic system takes on the role of expert biologists, reading each relevant published perturbation experiment, assessing the regulatory nature of the result and the strength of evidence, while considering the findings of other papers and the endpoint of the target trait. The agent’s goal is to produce an accurate network model of regulatory processes underpinning the trait. This model is deemed effective if it can faithfully reproduce the directional effects of known perturbations on the trait using signal propagation methods.

FLASH-P presents a major leap forward in mechanistic network creation from literature. Traditionally such literature-driven networks were manually constructed by domain experts - a painstaking process constrained heavily by time and resources^14^. These issues have been addressed by FLASH-P using a team of collaborative and competent AI agents to supercharge the perturbation curation and network synthesis process. Here we demonstrate that on-demand networks built by FLASH-P predict known perturbation outcomes with consistently high accuracy across a wide variety of traits in diverse plant species. Perturbation complexity ranges from single knockouts to multiplexed combinations of knockouts, over-expression, and environmental treatments. We further show that multiple single-trait networks can be intelligently merged by the agents, enabling multi-trait perturbation and pleiotropy analysis with minimal loss of model accuracy. This pleiotropy analysis is essential for prediction across multiple complex traits such as plant yield and response to stress.

## Results

### Overview of FLASH-P multi-Agent

FLASH-P is a multi-agent framework (Fig. 1b) that autonomously curates fragmented literature into perturbable, signed-directed causal network models. Given a target trait in a chosen species as input (Fig. 1a), it returns a network model of that trait in which nodes represent genes, metabolites and environmental factors, and edges represent the regulatory relationships that cascade to the target trait node. The effectiveness of the FLASH-P process is ensured by “Judge” agents that critique outputs at key checkpoints and allow progression only when defined metrics are met (Fig. 1c, methods). Perturbations are simulated on a network by setting modifier values on one or more nodes, followed by three different propagation methods (RWR, Algebraic, and ODE, methods). The final accuracy of a FLASH-P network model is determined by its ability to correctly recover the directional effects (up, down or unchanged) of known perturbations on the trait once propagation stabilizes.

**Fig. 1:**
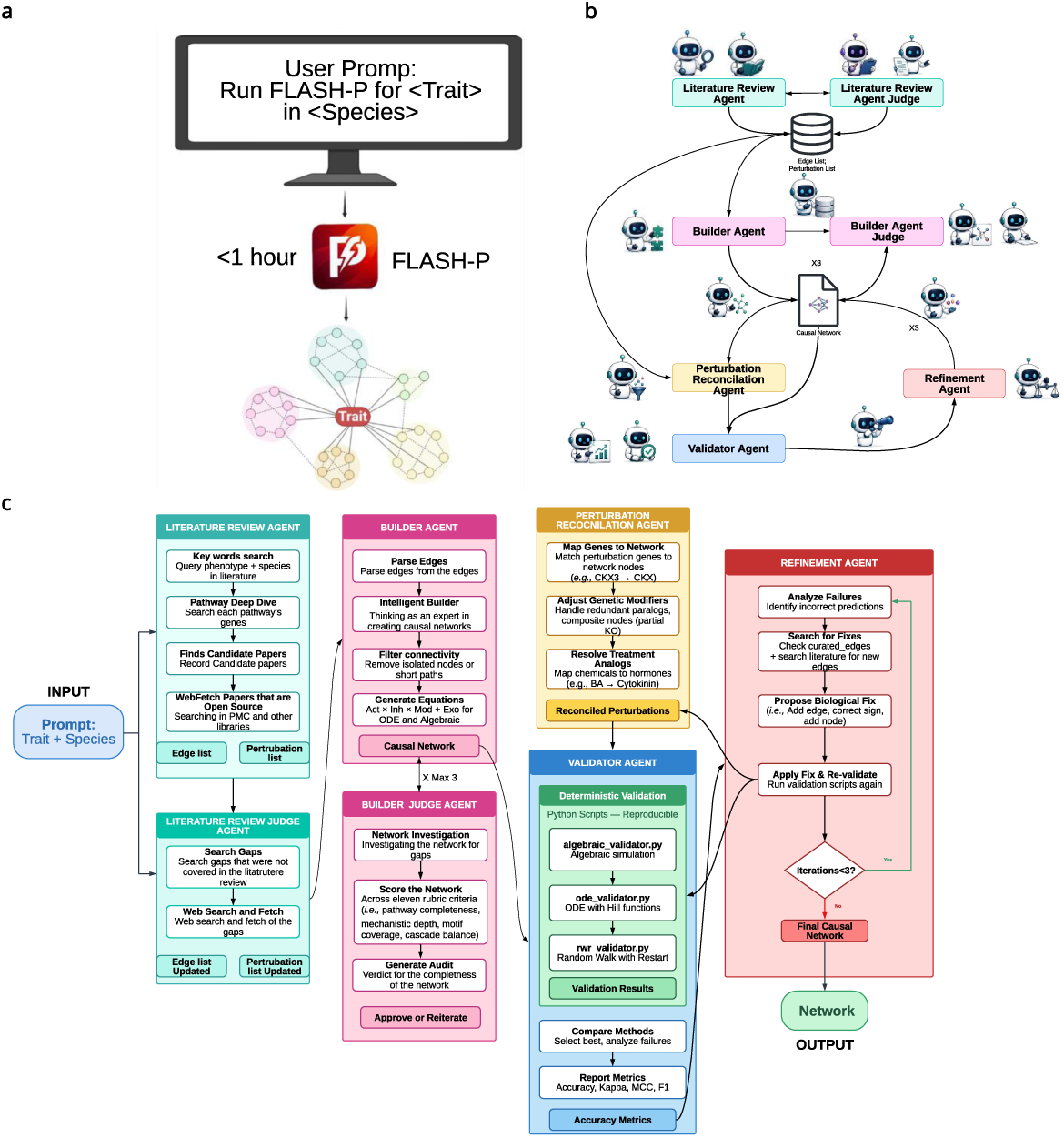
The FLASH-P pipeline for automated construction of causal regulatory networks from published literature. **a**, A single user prompt specifying a trait and species runs the full pipeline end-to-end and returns a validated causal network in under one hour. **b**, Pipeline architecture and information flow between the six agents. The Literature Review Agent and its independent Judge populate a shared repository of curated edges and perturbation experiments. The Builder Agent and its Judge assemble these edges into a causal network over up to three review cycles. The Perturbation Reconciliation Agent maps each literature experiment onto network nodes, and the Validator Agent simulates the perturbations and compares predicted phenotypes against reported outcomes. The Refinement Agent then revises the network in response to mispredictions, again over up to three iterations. Solid arrows denote data flow; dashed arrows denote control flow. **c**, Internal steps of each agent. From a trait–species input, the Literature Review Agent searches keywords, surveys pathways and fetches open-access full texts to build edge and perturbation lists; the Judge audits these and adds missing edges through targeted gap searches. The Builder parses edges, assembles cascades, prunes disconnected nodes and writes propagation equations under two frameworks (algebraic and normalised Hill ODE); the Judge enforces biological motifs and pathway completeness. The Reconciliation Agent matches gene names, assigns genotype modifiers and resolves chemical–hormone analogues. The Validator runs three deterministic Python simulators (algebraic propagation, Hill ODE, random walk with restart) and reports accuracy, Cohen’s κ, MCC and per-class F1. The Refinement Agent analyses failures, searches the literature for corrections backed by a published source and re-validates; the best-performing network across iterations is retained as output.

We used FLASH-P to autonomously construct and validate 12 single-trait networks, and a multi-trait merged network (AraMerged) comprising 6 single-trait Arabidopsis networks. Single-trait networks ranged from 35-80 nodes and 51-134 edges. Each network was generated in a zero-shot manner in less than an hour without significant computational power using Claude. A total of 1,088 published perturbation tests were curated by FLASH-P for validation of the single-trait networks, and an extra 35 for pleiotropy analysis on the AraMerged network.

### FLASH-P networks strongly predict known perturbation effects

The output of FLASH-P is a network model of trait regulation, and its accuracy is quantified by testing whether the network’s predicted perturbation responses follow the same direction as those reported in the literature. For the twelve single-trait networks FLASH-P achieved a mean best accuracy of 89.8%, where’best accuracy’ implies the highest accuracy amongst three network propagation methods (Table 1). The six Arabidopsis networks were particularly well constructed, with the shoot branching network reaching 96.8%. The framework also generalized well to biological systems beyond herbaceous plants, with both the *E. coli* lycopene production and the poplar lignin S/G ratio networks attaining high accuracy (Table 1). Overall, FLASH-P was consistent across diverse species and traits without requiring species-specific tuning or manual curation.

**Table 1.** Predictive accuracy of twelve FLASH-P-constructed single-trait causal networks against literature-reported perturbations. Each row reports a network built by FLASH-P for a given species–trait combination, listing the number of nodes and directed edges retained after construction, the number of perturbation tests reconciled from the literature (Tests), the highest classification accuracy obtained among the three network-propagation methods (Best Acc.), and the method that produced it (Best Method). The three methods are algebraic propagation, normalised Hill ODE, and random walk with restart (RWR). Accuracy is the fraction of perturbation experiments for which the predicted direction of phenotypic change at the trait node matches the experimentally reported outcome.

| Species | Trait | Nodes | Edges | Tests | Best Acc (%) | Best Method |
| --- | --- | --- | --- | --- | --- | --- |
| *A. thaliana* | Seed size | 80 | 134 | 102 | 91.2 | RWR |
| *A. thaliana* | Flowering time | 42 | 87 | 93 | 94.6 | Algebraic/RWR |
| *A. thaliana* | Lateral root density | 55 | 68 | 119 | 86.6 | All |
| *A. thaliana* | Shoot branching | 38 | 75 | 63 | 96.8 | ODE |
| *A. thaliana* | Hypocotyl length | 54 | 103 | 116 | 95.7 | Algebraic |
| *A. thaliana* | Plant height | 78 | 98 | 115 | 93.0 | RWR |
| *Z. mays* | Kernel row number | 50 | 80 | 71 | 85.9 | Algebraic |
| *S. bicolor* | Flowering time | 35 | 81 | 77 | 88.3 | Algebraic |
| *T. aestivum* | Plant height | 42 | 51 | 50 | 94.0 | ODE |
| *O. sativa* | Tillering | 69 | 100 | 165 | 80.0 | Algebraic |
| *E. coli* | Lycopene production | 62 | 92 | 86 | 84.9 | RWR |
| *Populus spp.* | Lignin S/G ratio | 44 | 96 | 31 | 87.1 | ODE |
|  |  |  |  | **1,088** | **89.8** |  |

To illustrate how a FLASH-P network models the regulatory processes underpinning a trait we trace a single gene knockout through the Arabidopsis shoot branching network (Fig. 2a,b,c). Under control conditions all nodes sit at a baseline value of 1 and the phenotype is at steady state (Fig. 2b). A knockout perturbation is introduced at the *MAX2* gene by setting its modifier to 0 (see Methods). This removes *MAX2*-mediated degradation, so its targets *SMXL6/7/8* and *BES1* accumulate and suppress *BRC1*, both directly and by repressing its co-activator *SPL9*. The loss of *BRC1* derepresses the auxin transporter *PIN3* and abolishes the *BRC1*→*HB21*→*NCED3*→ABA branch-inhibitory cascade, while *SMXL6/7/8* also up-regulates *PIN1*. Together, these effects predict an increase in shoot branching, matching the observed phenotype of *MAX2* KO mutants (Fig. 2c).

**Fig. 2:**
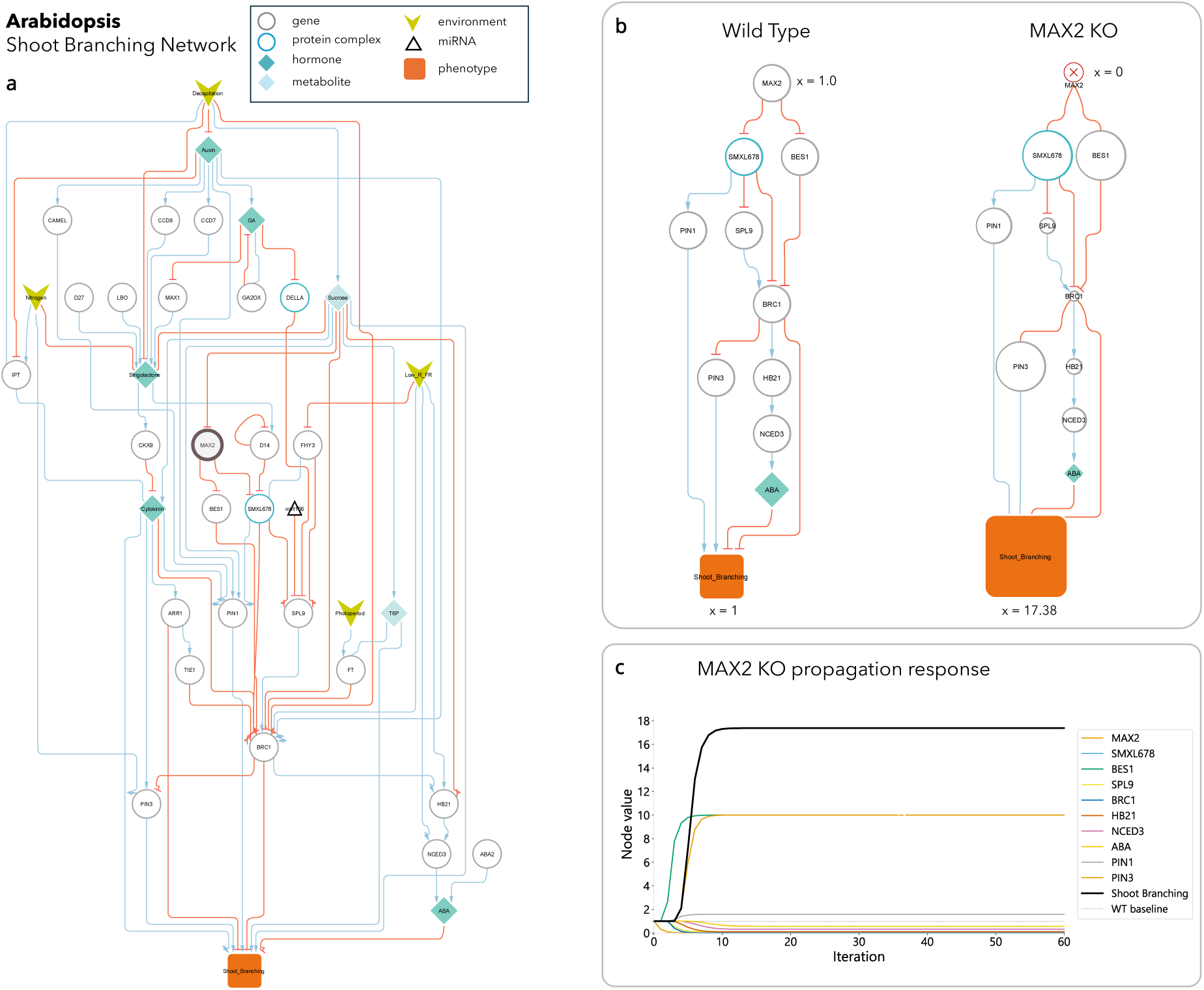
Perturbation simulation in the FLASH-P-constructed Arabidopsis shoot branching network. **a**, Full network produced by FLASH-P and visualized with Cytoscape^31^. Node shape and colour denote node class, edge colour distinguishes activation from inhibition. The MAX2 node, used as the perturbation example in **b** and **c**, is highlighted with a bold outline. **b**, Subnetwork view of the strigolactone signaling module under wild-type conditions (left) and under a MAX2 knockout (right; MAX2 modifier set to 0). Node sizes scale with the converged steady-state value x produced by the algebraic propagation framework. The shoot branching node value rises from x = 1 in wild-type to x = 17.38 in the knockout, recapitulating the increased branching phenotype reported for max2 loss-of-function mutants. **c**, Iteration-by-iteration trajectories of the nodes shown in **b** during propagation of the MAX2 knockout. Each trace gives the node value across 60 iterations of the algebraic propagation; the wild-type baseline of 1 is shown as reference. Trajectories converge to the steady-state values displayed in the right-hand network (not to scale) of **b**, illustrating how a single genetic perturbation propagates through the cascade to a directional phenotypic prediction.

### Agentically merged multi-trait network enables pleiotropic prediction

The AraMerged network preserved single-trait accuracy after integration, despite the presence of many more nodes, edges and alternate pathways between perturbations and their target traits. Accuracy across the 6 Arabidopsis traits dropped by only 1% (Fig. 3a). Importantly this demonstrates the ability to create much larger, yet still accurate, multi-trait network models. Furthermore, the AraMerged network supports pleiotropic prediction. We curated 35 pleiotropic perturbation tests covering 15 genes with documented effects on multiple traits. Known pleiotropic effects were recovered with high accuracy, usually by all three methods (Fig. 5a,b, green cells), with errors concentrated in Plant Height. These predictions arise from shared pathway nodes that connect traits through common signalling intermediates pathways. The cascade architecture in Fig. 4e is therefore not only the source of single-trait accuracy but also the structural prerequisite for pleiotropic prediction in an integrated network.

**Fig. 3:**
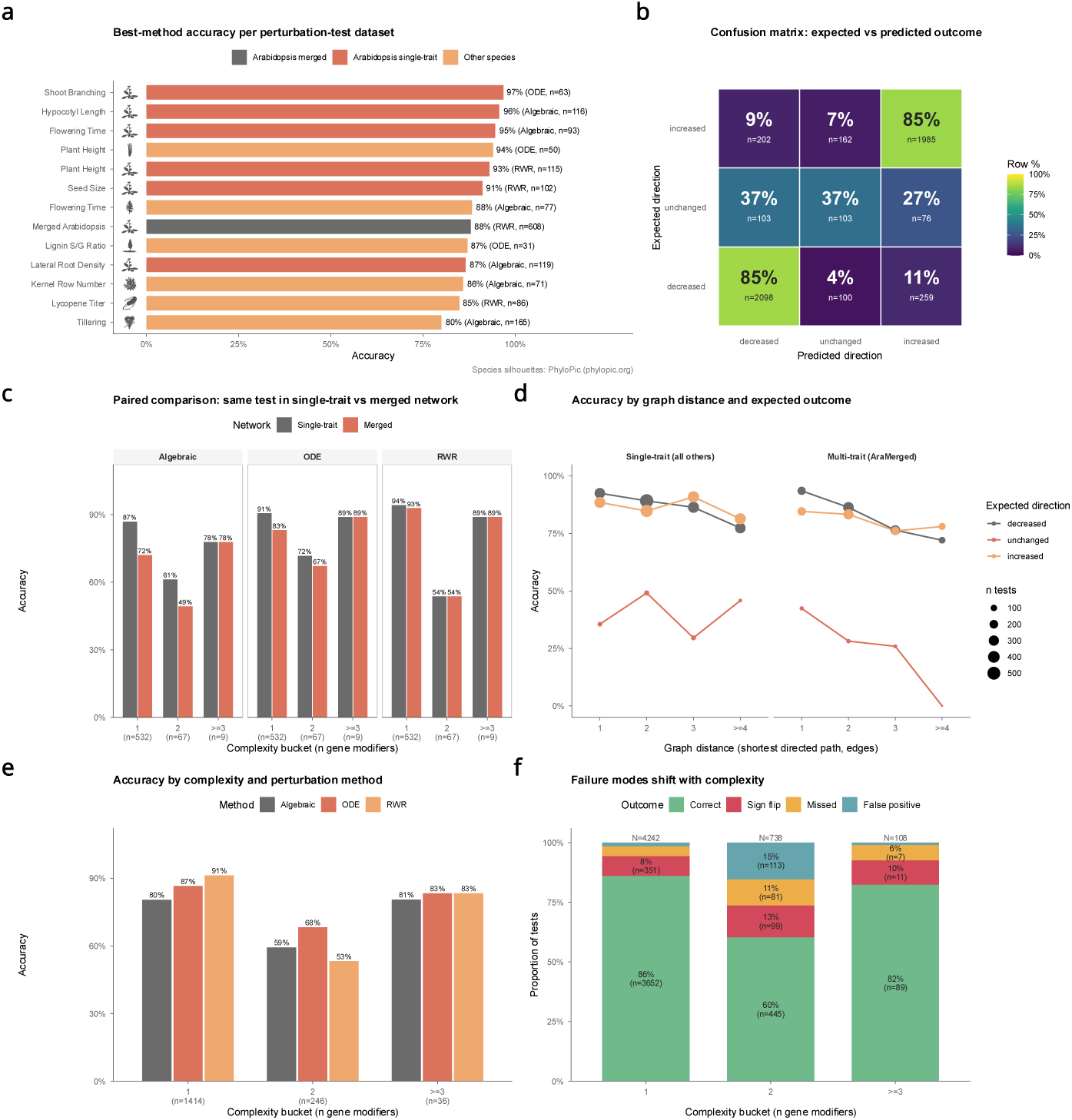
Factors affecting predictive accuracy in FLASH-P networks. **a**, Best-method accuracy for each of the twelve single-trait benchmarks and the merged Arabidopsis multi-trait network (AraMerged). Each bar is annotated with best accuracy, the propagation method that achieved it (Algebraic, ODE or RWR) and the number of perturbation tests *n*. **b**, Confusion matrix pooling perturbation tests across all networks and propagation methods. Rows give the experimentally reported (expected) direction of phenotypic change at the trait node; columns give the predicted direction. Cells show the row-normalised percentage and absolute count. **c**, Paired comparison of accuracy on the same perturbation tests run on the Arabidopsis single-trait networks versus the merged Arabidopsis network, faceted by propagation method and stratified by perturbation complexity (number of gene modifiers per perturbation: 1, 2 or ≥3). **d**, Accuracy as a function of the shortest directed path (in edges) between the perturbed node and the phenotype, separated by expected outcome. Left, all single-trait networks; right, the merged Arabidopsis network. Marker size encodes the number of tests at each distance. **e**, Accuracy by perturbation complexity bucket and propagation method, pooled across all networks. **f**, Failure-mode decomposition by complexity bucket: bars give the proportion of tests classified as correct, sign flip (opposite direction predicted), missed (no change predicted where a change was expected) or false positive (change predicted where none was expected). The merged Arabidopsis network is included alongside its constituent single-trait networks in c and d.

**Fig. 4:**
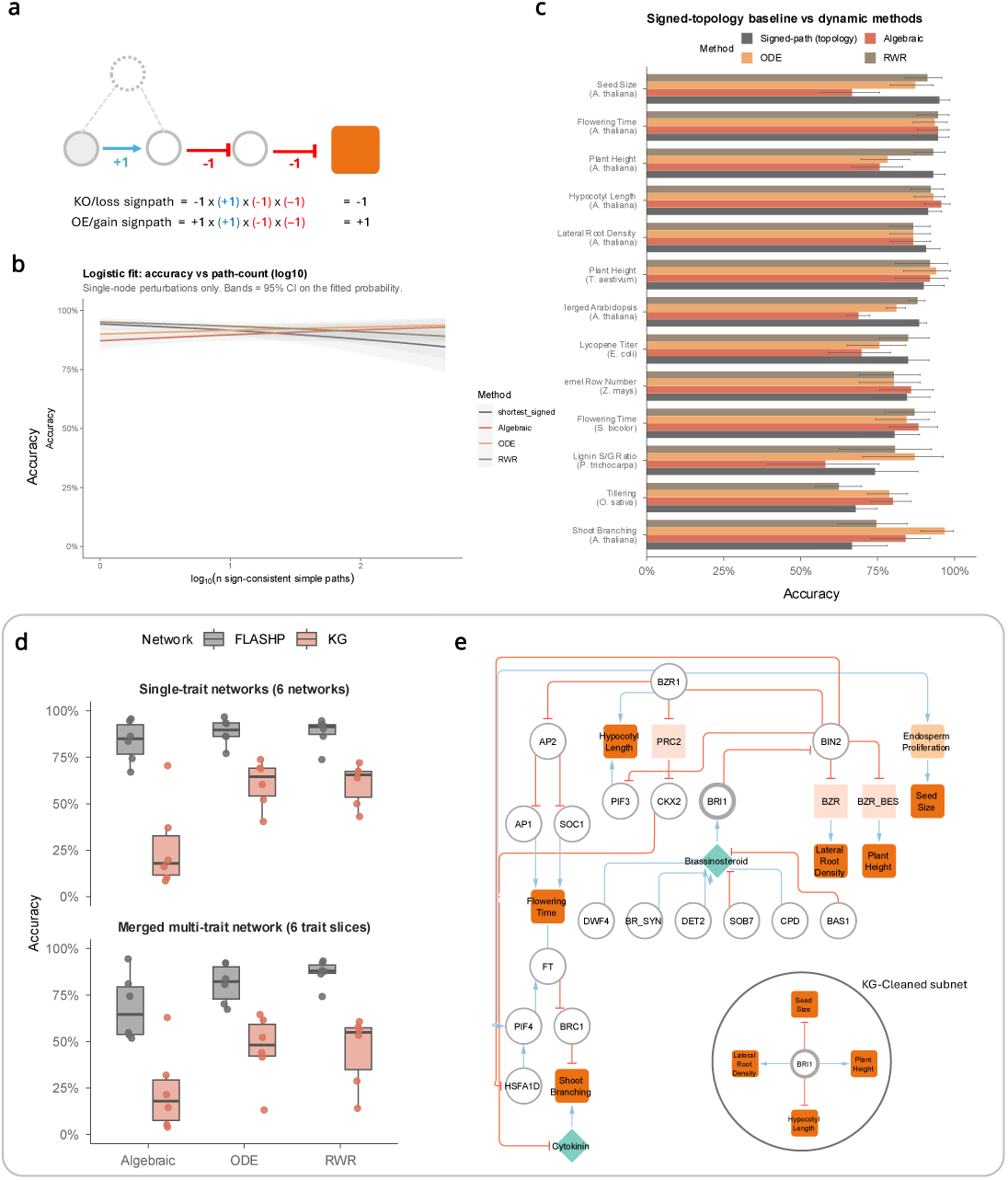
Network topology is the principal driver of predictive accuracy. **a**, Schematic of the signed-path method. The predicted direction of phenotypic change is the product of the perturbation sign (knockout/loss = −1, overexpression/gain = +1) and the edge signs (activation = +1, inhibition = −1) along the shortest directed path from the perturbed node to the phenotype. The method uses topology only, with no fitted parameters. **b**, Accuracy of the three propagation methods (Algebraic, ODE, RWR) stratified by whether propagation by network topology (signed-path) agrees with the experimentally reported outcome (left) or disagrees (right) **c**, Logistic fit of accuracy against the number of sign-consistent simple paths between the perturbed node and the phenotype (log10 scale), single-node perturbations only (n = 888). Bands, 95% confidence intervals. The signed-path baseline and RWR degrade as paths accumulate, while Algebraic and ODE remain robust by integrating signed contributions across alternative paths. **d**, Accuracy of FLASH-P networks versus networks built from a cleaned knowledge graph (KG) for the same six Arabidopsis traits, under each propagation method. Top, single-trait networks (one point per trait); bottom, the merged multi-trait network sliced by trait. Boxes, median and interquartile range; whiskers, 1.5× interquartile range. **e**, Topology around the brassinosteroid receptor *BRI1* in the merged Arabidopsis FLASH-P network (main diagram) and in the corresponding KG-derived network (inset, KG-cleaned subnet). In FLASH-P, *BRI1* propagates through cascade intermediates (*BIN2, BZR/BES1, PIF3, AP2, CKX2, PRC2*) before reaching phenotypes; in the KG, *BRI1* connects directly to each phenotype, leaving no intermediates through which a perturbation can propagate. Node shape and colour follow Fig. 2.

**Fig. 5:**
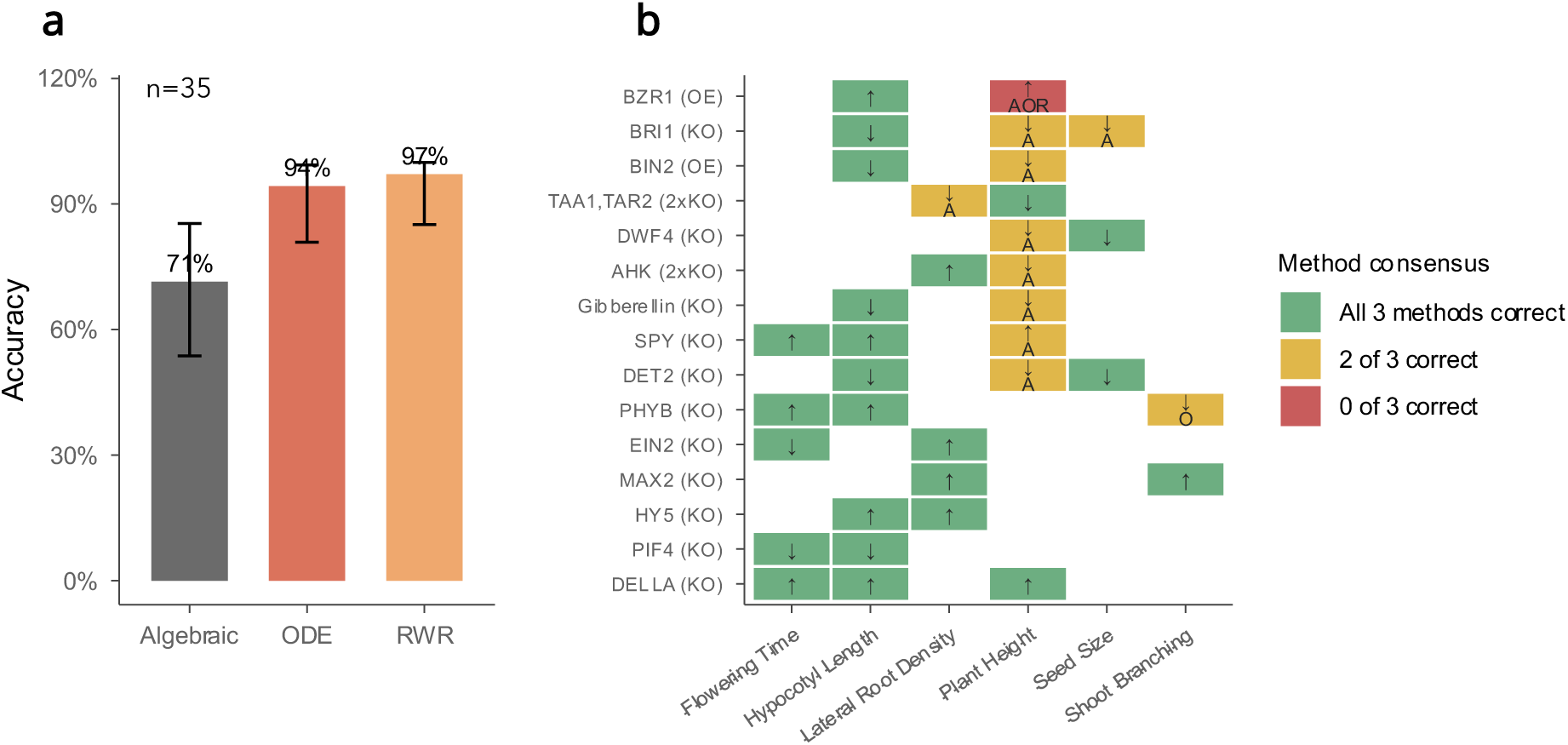
Pleiotropic prediction in the merged Arabidopsis multi-trait network. **a**. accuracy on a benchmark of 35 pleiotropic perturbation tests covering 15 genes with experimentally documented cross-trait effects, evaluated under the three propagation methods (Algebraic, ODE, RWR). Error bars show bootstrap 95% confidence intervals. **b**, per-test outcome matrix. Rows are perturbations (gene name and perturbation type: KO, knockout; OE, overexpression; 2xKO, double knockout) and columns are the affected phenotypes recorded in the literature. Cells are filled only where a literature-reported cross-trait effect exists. Cell colour indicates method consensus: green, all three methods predicted the correct direction of phenotypic change at the trait node; yellow, two of three methods correct; red, no method correct. Letters within yellow and red cells indicate which method(s) returned the incorrect prediction (A, Algebraic; O, ODE; R, RWR). Arrows give the experimentally reported direction of phenotypic change (↑, increased; ↓, decreased).

### Independent runs of FLASH-P generate variation for ensemble network creation

Given the non-deterministic nature of LLMs, we assessed the consistency of networks generated across independent runs by performing five additional runs for Arabidopsis shoot branching alongside the original. After ensuring all node identities were unified across the 6 shoot branching networks, we found that their topologies were substantially different, with 50% of the total edge pool being unique to a single network (Extended Data Fig. 1). Despite the differences, all six networks achieved comparably high accuracy, ranging from 85.9 to 97.1 % over 56 to 78 perturbations (Extended Data Table 1). Additionally, many perturbations could be tested in multiple branching networks due to the presence of identical nodes. These shared perturbations produced nearly 100 % accuracy in each network (Extended Data Fig. 2), indicating consistency of outcomes despite the topological variation.

Notably, every edge that was common to two or more networks had consistent sign across networks. This suggests that the topological variation between runs is driven by agents focusing on distinct parts of the underlying biology, not by hallucination. We leveraged this by using the merging agent to combine the six unique shoot branching networks into an ensemble network of 73 nodes and 177 edges, which we tested against 141 perturbations pooled across all runs, achieving 84.4 % accuracy (Extended Data Table 1). Most of the error stems from dilution of propagation signal across the larger graph rather than incorrect edges. The non-deterministic nature of the agents is therefore not a limitation but an asset, helping to construct the most comprehensive network for a given trait.

### Network and perturbation factors affecting prediction accuracy

To characterise where prediction errors arise, we jointly analyzed all perturbation results across the twelve single-trait and AraMerged networks.

Prediction errors concentrated predominantly on the *unchanged* class (Fig. 3b). *Unchanged* phenotypes were predicted correctly only 37% of the time while *increased* and *decreased* phenotypes were both predicted with 85% accuracy. This disparity follows from how signals move through a network. When a node is perturbed, the signal propagates along paths and either builds up or gets weakened along the way, but it almost always causes some impact at the trait node. Therefore, predicting an *unchanged* steady state is the hardest of the three classes. An *unchanged* outcome is also hard to verify biologically, since experiments are designed to detect differences, not to confirm their absence, so a real change can go unreported when replication is limited or conditions are not tightly controlled.

Accuracy for *increased* and *decreased* outcomes declined only mildly with greater network distance between perturbed and trait nodes, indicating that signal attenuation is not a limiting factor at path lengths typical of biological regulatory networks (Fig. 3d). *Unchanged* outcomes, by contrast, became substantially harder to predict as path distance grew.

With respect to perturbation complexity (Fig. 3e), single-modifier perturbations were predicted at 80 to 91% accuracy across the three methods, and triple or higher-order perturbations at 81 to 83%. Two-modifier tests, however, dipped to 58 to 66%. Failure-mode decomposition (Fig. 3f) localized this dip almost entirely to situations where an *unchanged* phenotype was expected but an *increased* or *decreased* phenotype was predicted (*i.e.* false positives). This pattern reflects how researchers usually pair two regulators that act in opposite directions, so their effects cancel at the phenotype. When the network propagates the two opposing signals, however, one exceeds the other and a directional change is predicted instead of the expected unchanged result. Note that here we compare each higher-order mutant against the wild type rather than against the corresponding lower-order mutant (for example, double versus single), but the difficulty remains, since the model struggles with unchanged outcomes for the reasons described above.

We next asked whether merging the six Arabidopsis single-trait networks into one (AraMerged) eroded accuracy for the same perturbations. Perturbation accuracy on the merged network remained high regardless of complexity, matching single-trait accuracy for high-complexity perturbations (Fig. 3c). The single-trait network model is therefore extensible to a multi-trait model via the Merge Agent.

### Prediction Accuracy is largely driven by topology

A perturbation propagates through a FLASH-P network along directed paths from the perturbed gene to the trait node, with each edge carrying a sign indicating whether the regulation is activating or inhibiting. The prediction made for a given perturbation could in principle depend on three things: the topology of the network (which genes connect to which, in what direction, and with what sign), the choice of propagation method that simulates how the signal travels along these paths, and any free parameters fitted within each method. Given the high accuracy achieved across all networks, we asked which of these three contributes most.

To represent the contribution of topology we introduced a measure in which the predicted direction of phenotypic change is simply the product of the initial perturbation sign and the edge signs along the shortest directed path (SDP) from the perturbed node to the trait node (Fig. 4a). Across most networks, the SDP performed comparably to the best dynamic method (Fig. 4b), indicating that the predictive signal lies primarily in the sign directed topology constructed by FLASH-P.

The propagation method does appear to matter under increased topological complexity between perturbation and trait (Fig. 4c). The complexity of the route between a perturbed gene and the trait is captured by the number of distinct directed paths connecting them. We tested how each method handled this complexity across 888 single-node perturbation tests, pooled across all twelve networks, using logistic regression of correctness on path count (Methods)**Fig. 4**. When only one possible signed path existed, the SDP reached 98% accuracy. With multiple competing paths, the SDP accuracy declined significantly (*P* = 0.016; logistic regression; Methods) as did random walk with restart (RWR) (interaction *P* = 0.74), since both propagate along edge signs without integrating information across paths. Algebraic and ODE, on the other hand, maintained 88 to 91% accuracy at over ten competing paths, declining significantly less steeply (interaction *P* = 0.006 and *P* = 0.015 respectively). FLASH-P networks therefore retain accuracy in topologically complex regions because Algebraic and ODE methods can integrate contributions from multiple competing paths.

### FLASH-P models the regulatory processes underpinning the trait, which are required for accurate predictions

FLASH-P networks are generated very differently to knowledge graphs (KGs) which aggregate published gene-to-trait associations and interactions between genes and other factors. Nevertheless, we evaluated whether curated trait-specific subnetworks (KG-Cleaned) extracted from the PlantConnectome KG could act as a trait model in the same manner as FLASH-P networks by running matched perturbation simulations on both.

FLASH-P networks strongly outperformed KG-Cleaned networks for every Arabidopsis trait and propagation method (Fig. 4d). This result is due to the nature of KGs where any gene-trait association is wired as a direct path to the trait (pathlength=1), even if the true regulatory mechanism requires a complex cascade of intermediates. KG-Cleaned therefore performed best on traits where many nodes happen to be true direct regulators of the trait, such as Shoot Branching (73.8% accuracy), but poorly for traits where many regulators operate through intermediary pathways. FLASH-P, on the other hand, maintains accuracy because it is designed to explicitly model these pathways. The performance gap between the two therefore measures how much of the accuracy depends on the network carrying the intermediate regulatory steps rather than a list of direct gene-to-trait edges.

In AraMerged, FLASH-P maintained 69.6 to 88.5% accuracy across the three methods on 608 single-trait tests and 71 to 97% on 35 pleiotropic outcomes, whereas equivalent merged KG-Cleaned network dropped to 20.1 to 42.9% and 20.0 to 48.6% respectively. In the merged KG-Cleaned network each gene remains connected only to its originally catalogued trait, so a perturbation for one trait cannot reach another trait node, thus pleiotropic effects become undetectable.

## Discussion

Decades of individual perturbation experiments remain fragmented across thousands of papers with inconsistent formats and varying genetic backgrounds, which limits our ability to use it for purposes beyond simple knowledge aggregation. FLASH-P demonstrates that agentic AI can unlock new capabilities from this fragmented knowledge with a very high level of efficiency and accuracy. We have shown that FLASH-P synthesises scattered literature into predictive causal networks from a single prompt, recovering the direction of 1,088 published perturbations with a mean accuracy of 89.8% across twelve networks spanning seven species. Its real power, however, lies in the Merge agent, which intelligently combines several single-trait networks into one. For example, the Merge agent can take advantage of the stochastic nature of the underlying LLM by consolidating independent runs of the same trait into a more comprehensive ensemble network. Multiple runs of FLASH-P produces different models of the same trait. Each model is a good fit to the existing perturbation knowledge, and no single model can be considered “correct”. This effectively emulates the output of a team of experts, where each is likely to produce a good result while investigating certain aspects of the system more deeply than others. More impressive still is the ability to merge multiple networks of different traits into one multi-trait network that recovered known pleiotropic effects that no single network could capture. These results highlight the potential of the approach to scale the creation of mechanistic models underpinning any trait of interest, as a single network, an integrated multi-trait model, or a consolidated single-trait one.

The prevailing approach for utilising literature-based knowledge is the aggregated knowledge graph. Recently, KGs have been efficiently produced by AI-automated extraction of entity-entity relationships from online resources^15–17^, providing a massive representation of known biological interactions. A knowledge graph represents the interaction space directly, storing relationships as a static inventory of gene-to-trait associations. FLASH-P networks are also built through AI-automated extraction of known interactions, but they differ in agentic reasoning and intent. FLASH-P uses the regulatory relationships among all nodes to build the layered intermediate steps through which perturbation signals propagate to the trait. Its perturbation accuracy therefore comes from the hierarchical gene-to-trait causal topology itself, agreeing with Santolini and Barabási^18^, who showed that signed directed topology is a primary determinant of predictive power, recovering up to 80% of perturbation patterns even without kinetic parameters.

The ability to automatically construct predictive causal networks models has practical significance only if these networks can be applied usefully in downstream applications. We highlight several potential areas in plant breeding, gene editing and fundamental plant science.

In plant breeding, Wallace^13^ discussed the transition from Breeding 3.0, where genome-wide markers enable statistical prediction of breeding values, to Breeding 4.0, where breeders can identify the optimal combination of alleles for a target environment and stack them into ideotypes via genomic selection or gene editing. Research is working toward this goal through complementary approaches, including the integration of crop growth models with whole-genome prediction to capture genotype-by-environment interactions ^19–21^, and the development of hierarchical gene-to-phenotype maps that decompose complex traits into intermediate physiological components^22^. Constructing the mechanistic networks that connect genetic variation to physiological parameters has historically required extensive manual curation and domain expertise, making it impractical at the scale needed across traits and species. FLASH-P removes this bottleneck by generating a validated causal network for any trait on demand, making it feasible to build the mechanistic layer that crop growth models such as APSIM require to link genotype to physiological parameters^23–25^.

FLASH-P networks could boost the accuracy of genomic prediction models by enabling biologically informed selection and weighting of causally relevant loci. Farooq^26^ demonstrated in Arabidopsis that grouping markers by Gene Ontology terms and gene co-expression networks improved prediction of growth-related traits, and noted that heterogeneous sources such as interaction networks and pathway databases could be combined to pinpoint candidate QTLs. FLASH-P networks could serve as improved biological priors in genomic prediction models like BayesRC^27^, informing both marker class assignment and the expected directionality of marker effects based on their position in the regulatory cascade. Merged multi-trait networks could likewise inform multi-trait genomic prediction models that account for genetic correlations through documented biological mechanisms rather than purely statistical covariance. Such multi-trait networks could also be used to prioritise selection strategies based on a predicted likelihood of having a negative consequence on non-focal traits or highlight strategies to retain important genetic diversity in the network.

The networks also could have practical value for experimental design, since perturbation-informed methods can prioritise the most informative targets and reduce the number of costly experiments needed to identify regulatory interactions^28^. This remains a hypothesis, however, as the networks have not yet been tested on novel perturbations. We leave this validation to future work, since it falls outside the present scope of developing the multi-agent pipeline. The speed of network construction also lends educational value, offering a structured way to organize large bodies of biological knowledge and reveal modules and pathways that would be difficult to assemble manually.

Several limitations should be acknowledged. First, paywalled publications may contain important perturbation data but are currently inaccessible to the Literature agent. As agentic AI becomes more prevalent in scientific research, structured machine-readable access to the full literature would substantially improve automated knowledge extraction. We did not connect the current agent to structured plant resources because perturbation experiments with phenotype outcomes are not yet curated at sufficient scale in such databases^29,30^, but future versions could integrate them as complementary retrievers. Second, the perturbation tests used for validation primarily involved single-gene knockouts and simple treatments, since these dominate the literature. Double and triple-or-higher perturbations were tested against wild-type, but we did not test epistasis, which requires comparing double mutants against their constituent single mutants, something that could be readily addressed by a single change in the Agent’s instructions from the user. Third, an inherent possibility of LLM hallucination remains. An automated audit (Supplementary Note 1) found that 98.8% of edges and all perturbations logged by FLASH-P carried a valid DOI, with 2.4% citing a DOI that did not resolve, and a verification sample of 96 citations found 92.7% substantiated their claim. These are hallucinations in the DOI itself, not wrong edges, and a genuinely wrong edge would have been flagged and removed or corrected by the judges in the FLASH-P pipeline. This is why the six shoot branching networks contain no contradicting edges, but 100% agreement, and any possible initial errors that may happened were caught by the validator agents (Extended Data Fig. 1). Finally, we validated networks against perturbation experiments already documented in the literature. Whether they can accurately predict the outcomes of novel, untested perturbations lie outside the scope of this work, which focused on showing that large causal network models can be constructed rapidly and predict vast and diverse existing experimental knowledge.

## Methods

We developed FLASH-P, a multi-agent pipeline that autonomously assembles and validates directed, signed causal networks from published literature (Fig.5). When FLASH-P is prompted with “Run the full pipeline of FLASH-P for [trait] in [species]”, the agentic team proceeds by searching relevant literature, identifying molecular components, assembling the network, and testing whether the resulting network could accurately predict the phenotypic outcomes of perturbation experiments reported in the literature (Fig 5a,b,c). The network is finally refined by the agents up to three times to maximise predictive accuracy. The reasoning and analysis is performed in stages by established large language models (LLMs) as well as custom FLASH-P scripts, directed by the agentic team.

### Multi-Agent Architecture

#### Literature Review agent

The Literature Review agent identifies all documented regulatory interactions and perturbation experiments relevant to the specified trait and species (Fig.1B). The agent conducts at least ten rounds of iterative searches across multiple academic databases and publisher repositories using keywords covering hormones, transcription factors, environmental cues, and mutant phenotypes. Candidate papers are read by sub-agents that fetch full-text articles from open-access sources, extracting information from gene knockouts, overexpression studies, chemical treatments, and epistasis analyses. When open-access is unavailable, the agents fall back to abstracts. Crucially, the agents extract complete signaling cascades with all intermediate nodes, capturing, for example, the full sequence from hormone biosynthesis through receptor binding, signal transduction, and transcription factor activation rather than collapsing these into a single process. After compilation, the agent reviews the node and edge list as a domain expert would, identifying gaps in pathway coverage or missing feedback loops and running targeted follow-up searches to fill them. All extracted nodes and edges are supported by literature evidence and metadata, producing the curated edge set and raw perturbation dataset both of which are transparent to the user and can be manually reviewed if desired.

#### Literature Review Judge agent

The Literature Review Judge agent is an independent expert auditor that examines the Literature Review agent’s output before network construction begins. Acting as a senior domain expert with fresh eyes, it first enumerates from its own pre-trained field knowledge the canonical pathways, hub regulators, mots common mutants, environmental inputs and crosstalk loops expected for the phenotype, writing this checklist down before inspecting the extracted repository. In this way the audit is not shaped by what was already found by the Literature Review agent. It then systematically compares this canonical picture against the candidate papers, curated edges and perturbation dataset, flagging missing pathway arms, under-represented master transcription factors, receptors whose ligands are present but whose receptor is absent, one-sided bidirectional coverage where upstream regulators were overlooked, absent canonical mutants, and full-text papers that yielded far fewer edges than expected (for example a review paper on the phenotype that produced only one or two edges). For every issue classified as high or medium severity, Literature Review Judge agent runs targeted second-round searches across academic databases and publisher repositories, reads the newly discovered papers using the same extraction standards as the original review, and appends the new papers, edges and perturbation tests to the existing files with sequentially continuing identifiers. An untouched snapshot of the original output is preserved before any changes are made. This is to be transparent and the user can know exactly what has been done in each step. The agent finally produces a detailed audit report documenting every gap identified, every search performed, what was closed and what remains unaddressed due to paywalls or genuine literature limits.

#### Builder agent

The Builder agent transforms the curated edges into a causal network with activation or inhibition relationships between nodes. To make the network predictive rather than merely descriptive, the agent defines an information propagation system (Equation 1) where the value of each node at iteration *t* is computed from the values of its regulators at iteration *t-1*.

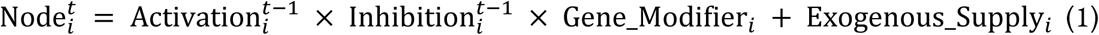

Node modifiers (value) provide a mechanism for simulating perturbations and examples values of nodes such as knockouts (modifier = 0), knockdowns (modifier = 0.5), wild-type baselines (modifier = 1), and overexpression (modifier = 2). The exogenous supply provides capacity for external inputs such as hormone treatments or environmental factors like light and nutrients. Under control conditions, with all genotype modifiers set to 1 and no exogenous supply, every node converges to a baseline value of 1, representing the unperturbed steady state. This provides a key benefit to the approach which is to enable a network at steady state to easily scale in size and that all perturbations are relative to this steady state enabling disparate experimental data to be utilized. The activation and inhibition terms in *eq1* are defined by two alternate mathematical frameworks: Algebraic and normalized Hill ODE.

In the algebraic framework, the activation term is the geometric mean of all activating inputs:

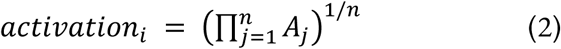

where *A*_*j*_ is the value of the *j*-th activator at *t-1*. The inhibition term uses a bounded inverse function where the product of all inhibitor values is inverted with a regularisation constant preventing division by zero and a cap on maximum de-repression.

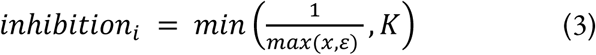

where 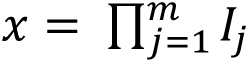 is the product of all inhibitor values at *t-1, ε* is a regularisation constant set to 0.1 to prevent division by zero, and *K* = 10 to cap de-repression.

In the normalized Hill ODE framework, activation and inhibition are instead defined using normalised Hill functions:

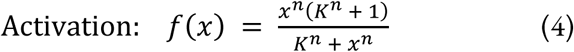

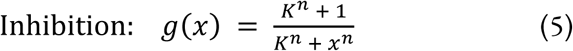

where *x* is the input signal (activator or inhibitor value), *n* is the Hill coefficient controlling switch-like behaviour and *K* is the half-maximal activation constant, providing a smooth sigmoidal response to input changes.

Both frameworks produce identical predictions under wild-type conditions and are tested against the same perturbation dataset, allowing direct comparison of their predictive performance. Rather than constructing a flat knowledge graph where all reported regulators connect directly to their target trait, the builder agent organises edges into layered signal pathways where perturbations propagate predictably through intermediate nodes to the phenotype. The agent does this using instructions and examples held in its.md files, together with LLM reasoning. This distinction is important because direct connections from many regulators to a single target node dilute individual perturbation signals through the geometric mean, making the network descriptive of the literature but unable to predict experimental outcomes. The agent enforces regulatory structure by limiting the number of activators and inhibitors per node, routing redundant regulators through pathway intermediates, and removing dead-end nodes that cannot influence the phenotype.

### Builder Judge agent

The Builder Judge agent conducts an independent biological quality review of the network after it has been built but before any perturbation test is run. Approaching the network the way a peer reviewer would approach a methods figure, the agent reads every node and every edge, cross-references the network against the full curated edge repository to identify which edges were rejected, and systematically scores the network across eleven rubric criteria: pathway completeness, mechanistic depth, motif coverage, cascade balance, composite node handling, hub completeness, topology hazards, evidence quality, phenotype audit, rejected edges review and key player density. For every node the agent produces a short audit listing its inputs, its outputs, any curated regulators that were left out, and a verdict on whether the node is well connected, is missing known regulators, or carrying more connections than the literature supports.

For every major pathway the agent walks the cascade end-to-end from source to phenotype, verifying that the network topology reflects the underlying biology rather than a flattened list of interactions.

As a quantitative check, the per-node coverage ratio compares the Literature Review agent’s curated edges against edges retained in the assembled network. Any node that had five or more curated edges but fewer than 30% (retention < 0.30) of these are retained in the Builder’s network is flagged as an under-represented hub. This guards against the Builder agent dropping too many edges during construction of the network, which would otherwise make a major regulator look like a minor one and bias the perturbation results. Crucially, the Builder Judge agent never reads the perturbation dataset, because its job is to assess biological credibility alone rather than predictive accuracy. It suggests rather than edits, writing a structured review with prioritised actionable recommendations tied to specific edges. The Builder then applies those suggestions and the review cycle repeats until the Judge agent returns a verdict of “approved” or a hard cap of three iterations is reached.

### Perturbation Reconciliation agent

The Perturbation Reconciliation agent is responsible for determining which of the perturbation experiments extracted by the Literature Review agent can be simulated on the constructed network. To do so, it must match literature gene and molecule names to network node names, exclude any perturbations for nodes removed during network building, and assign the appropriate modifier values and exogenous supply terms for each testable experiment. The agent also selects the appropriate baseline for each perturbation. Not all perturbations are compared against wild-type. A mutant treated with an exogenous compound, for instance, is compared against the untreated mutant, isolating the effect of the treatment from the genetic background.

### Validator agent

The Validator agent simulates each curated perturbation on the network using one of the three propagation methods described below, and compares the resulting phenotype value to a baseline value taken from a matching reference simulation (Fig. 1B). The choice of reference depends on the experiment type. For single-gene mutants and standalone treatments, the reference is the wild-type simulation, in which every node converges to 1 under control conditions, so the baseline phenotype value is 1. For rescue experiments, in which an exogenous compound or environmental cue is applied to a mutant background, the reference is the simulation of the mutant alone, and the baseline is the converged phenotype value of that mutant simulation. Because the baseline can be 1 or any other value depending on the test, the perturbed phenotype is divided by its baseline to produce a ratio that is comparable across all experiment types. A ratio of 1 indicates no change relative to the baseline, a ratio above 1 indicates a phenotype increase, and a ratio below 1 indicates a phenotype decrease. The direction of change at the phenotype node is then classified as *increased*, *decreased* or *unchanged* using a symmetric 5% tolerance band around 1:

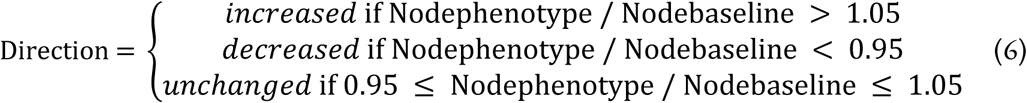

where Node_phenotype_ is the converged steady-state value of the phenotype node under the perturbation, and Node_baseline_ is its value in the matching reference simulation.

Three propagation methods are implemented and tested against the same perturbation dataset: algebraic propagation, normalised Hill ODE, and random walk with restart (RWR). All three use synchronous (Jacobi) updates, meaning that at every iteration the new value of each node is computed from the previous iteration’s values of its regulators, and all nodes are updated together. The loop terminates when the maximum absolute change across nodes falls below a small tolerance:

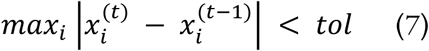

where *x_i_^(t)^* is the value of node *i* at iteration *t*, and tol = 1×10^−4^ (0.0001). A hard iteration cap stops the loop if convergence is not reached: 100 iterations for the algebraic method, 500 steps at d*t* = 0.1 for the ODE method, and 50 iterations for RWR.

#### Algebraic propagation

The algebraic framework applies the perturbation through the node modifiers and exogenous supply terms, then iteratively evaluates the master equation (Equation 1) for each node until convergence. To prevent oscillations that can arise from synchronous updates, the new value is blended with the previous value through a damping factor:

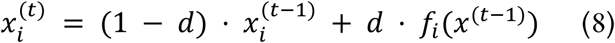

where *d* is the damping factor (a fraction of the newly computed value blended with 1 − *d* of the previous value), and *f_i_*(*x*^(*t*−1)^) denotes evaluation of the master equation (Equation 1) on the regulators of node *i*.

#### Normalised Hill ODE

The ODE framework models the network as a continuous dynamical system and solves for steady states numerically. It replaces the algebraic activation and inhibition terms with the normalised Hill kernels defined in Equations 4 and 5 (normalised so that *f*(1) = *g*(1) = 1, ensuring that the wild-type baseline is preserved) and integrates a first-order ordinary differential equation per node, with unit decay rate, using forward Euler:

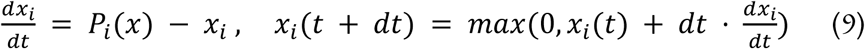

where *P*_*i*_(*x*) is the production term, computed from the master equation (Equation 1) as the product of the Hill activation and Hill inhibition evaluated on the regulators of node *i*, multiplied by the gene modifier and added to the exogenous supply. The skeleton is identical to the algebraic *A* · *I* · *g* + *e*, but with Hill kinetics in place of the geometric mean and bounded inverse.

Two parameters of the Hill kernels, the half-maximal activation constant *K* and the Hill coefficient *n*, are varied jointly to identify the optimal configuration for each network. All combinations of *K* ∈ {0.1, 0.5, 1.0, 2.0, 5.0, 10.0} and *n* ∈ {1, 2, 3, 4} are tested. All cells use identical d*t* = 0.1, max_time = 50, and convergence tolerance = 0.0001.

#### Random walk with restart (RWR)

The RWR framework propagates a signed perturbation signal rather than an absolute activity level. Each node starts with a restart value *S_i_*^(0)^ derived from its gene modifier: −1 for knockout, −0.5 for knockdown, 0 for wild-type, and +1 for overexpression. At every iteration the signal is updated as a convex combination of the mean signed signal from the node’s regulators and the restart vector:

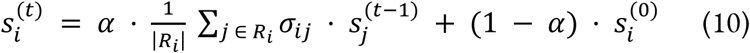

where α is the restart probability that controls how far the signal travels before being pulled back to its source, *R*_*i*_ is the set of regulators of node *i*, |*R*_*i*_| is its cardinality, and σ_*ij*_ ∈ {+1, −1} is the sign of the edge from *j* to *i*.

After convergence, each node is classified as *increased*, *decreased* or *unchanged* using a small direction threshold of 0.00001: signals above the threshold are reported as *increased*, below the negative threshold as *decreased*, and within the band as *unchanged*. As in the ODE framework, α is selected by a sensitivity sweep, here over {0.5, 0.6, 0.7, 0.75, 0.8, 0.85, 0.9, 0.95, 0.99}.

For rescue experiments, in which an exogenous compound is applied to a mutant background, the RWR system is run twice, once for the mutant alone and once for the mutant with the treatment, and the predicted direction is defined by the difference between the two phenotype signals.

The signed RWR method has no inhibitory production term, so a perfectly cancelling pair of signals is required to predict an *unchanged* outcome at the phenotype. In practice this rarely occurs, which is why the *unchanged* class is the hardest to predict of the three for this method.

#### Metrics and failure analysis

For each method the Validator reports overall accuracy, Cohen’s kappa, Matthews correlation coefficient and per-class F1 scores, together with bespoke scores that account for network complexity and for the number and complexity of perturbations (Supplementary Note 2). Because validation must be deterministic and reproducible, it is implemented as standalone Python scripts rather than delegated to the LLM. A failure analysis then examines each incorrect prediction to identify potential missing edges or mis-specified regulatory relationships, which informs the subsequent refinement stage.

#### Refinement agent

Following initial validation, a Refinement agent iteratively improves network accuracy by analysing incorrect predictions and searching the literature for biological explanations for the failure (Fig.1B). The agent proceeds to apply network changes that are backed by a published source with a verified DOI. The refinement process runs for a maximum of three iterations. At each iteration the modified network is re-validated using all three validation methods, and the iteration’s refinement is only accepted if it improves perturbation accuracy. The best-performing network across all iterations is selected as the final model.

All agent files can be found in https://github.com/CMits/FlashP.git.

#### Agentic Choices

FLASH-P decomposes network construction into seven specialised agents rather than a single generalist, because role-specific prompts and tool surfaces produce more reliable behaviour than one agent reasoning over the union of all tools. Each agent is restricted to the operations its role requires. Each agent’s prompt further decomposes its task into an ordered sequence of sub-steps and instructs the agent to reason through each sub-step before producing its output, which constrains the action space at each turn and reduces the variance of agent behaviour across runs. Stages communicate through curated handoffs rather than a shared transcript, so each agent receives the structured artefact it needs from the previous stage without inheriting the full upstream reasoning trace, which keeps LLM context windows tractable and prevents noise from earlier steps propagating forward.

All agent outputs are validated against a standard schema to prevent the generative nature of the agents from producing inconsistently structured files. Outputs that fail validation are returned to their originating agent with the schema error appended to context, with up to three retries before the run is flagged for review. To handle cases where a single pass misses relevant information, we introduced Judge agents in the Literature Review and Builder stages, creating an internal reason-and-check loop within each stage before its output propagates downstream. When a judge rejects an output, the originating agent is re-invoked with the judge’s critique in context; the loop terminates when the judge accepts the output or after three iterations.

Every edge in the final network retains provenance back to the literature source, passage and agent decision that introduced it, so the constructed network is fully auditable and supports the expert-feedback curation workflow.

#### Single-trait networks

To evaluate the generality and predictive accuracy of FLASH-P, we constructed causal regulatory networks for twelve single traits spanning seven species. Six networks were built for *Arabidopsis thaliana*: shoot branching, flowering time, hypocotyl length, plant height, lateral root density, and seed weight. Four networks were for agronomically important traits in crop species: rice tillering, sorghum flowering time, maize kernel row number, and wheat plant height, all of which are central to breeding programmes because they directly influence yield potential, harvest timing, and crop architecture under field conditions. Finally, a network for lignin S:G ratio was constructed for the model tree species Poplar, and a network for lycopene production was constructed for the prokaryote *Escherichia coli* to test whether the causal cascade framework generalises beyond herbaceous plants. Traits were selected to represent phenotypes that are quantitatively measurable and regulated by well-characterised but sufficiently complex pathways, deliberately avoiding highly integrative traits such as yield or disease resistance where hierarchical ontology-based decomposition would likely be required before causal modelling becomes tractable.

For each network, FLASH-P assessed its predictive accuracy by simulating literature-based perturbations and observing the effect on the phenotype node. A prediction was scored as correct when the simulated direction of change at the phenotype node matched the experimentally reported outcome.

#### Multi-trait merged networks

Biological pathways do not operate in isolation, so we developed an additional agent to merge single-trait networks into a unified multi-trait model. This is important because crop breeders are often interested in pleiotropic effects across traits and gene editing is often subject to off-target effects. The agent is necessary because a simple union of the single-trait networks would not produce a functional model for multiple reasons: each single-trait network was parameterized for predicting its specific target phenotype, not the other ones; shared nodes may carry different names across networks; common edges may conflict in sign; and the cascade structure must be reorganised at a larger scale to maintain signal propagation integrity. The Merge agent addresses this by first normalising node names across networks to unify shared components, then taking the union of all edges and resolving conflicts where the same regulatory interaction appears with contradictory signs by retaining the direction supported by more independent literature sources. The merged network is then filtered for connectivity, and equations (algebraic and ODE) are regenerated to reflect the new topology. All perturbation tests from the individual networks are pooled and validated using the same three propagation methods applied to individual networks, allowing direct comparison of per-trait accuracy before and after merging.

For this study we used FLASH-P to merge all 6 of the single-trait Arabidopsis networks into one multi-trait network (AraMerged). For validation, in addition to the pooled perturbations from the single-trait networks, we searched the literature for experiments that explicitly tested pleiotropic effects across the merged traits in the same species, providing an independent assessment of whether the merged network can predict cross-trait outcomes that no single-trait network could capture.

#### Comparison with knowledge-graph perturbations

To assess whether the cascade architecture constructed by the Builder agent improves predictive accuracy, we compared FLASH-P networks against flat knowledge graphs (KGs) extracted from PlantConnectome^15^. This is a state-of-the-art plant KG containing over 4.7m relationships parsed by LLM from over 71,000 papers. We queried PlantConnectome for each of the six Arabidopsis traits using seed terms comprising key pathway genes, hormones, and phenotype descriptors relevant to each trait, with the complete list of query terms provided in Supplementary Notes 3. Both network types were validated against the same set of perturbation experiments

The raw extracted KGs contained substantial noise including contradictory edge signs, redundant entries, free-text node names with multiple gene identifiers, and edges with unknown regulatory direction, making direct simulation infeasible. We therefore applied a cleaning script that retained only Arabidopsis-annotated edges, removed edges with unknown direction, extracted and normalised gene names to a common nomenclature, and resolved conflicting edge signs by retaining the direction supported by more literature sources. We called these networks KG-cleaned. Algebraic and ODE equations were then auto-generated directly from the resulting network topology using the same mathematical frameworks applied to FLASH-P networks.

#### Statistical test for the effects of network complexity

To test whether the complexity of the network between the perturbed node and the trait node has an effect on accuracy, we introduced a measure in which the predicted direction of phenotypic change is simply the product of the initial perturbation sign and the edge signs along the shortest directed path (SDP) from the perturbed node to the trait node. We evaluated the propagation methods against an SDP baseline using a logistic regression fitted on single-node perturbations pooled across the twelve single-trait networks (*n* = 888 tests). Per-test correctness was modelled as a function of the log-transformed number of sign-consistent simple paths from the perturbed gene to the phenotype node, the prediction method, and their interaction, with the signed shortest-path baseline taken as the reference method. For test *i*, let *y*_*i*_ ∈ {0,1} denote whether the prediction was correct and let *p*_*i*_ =

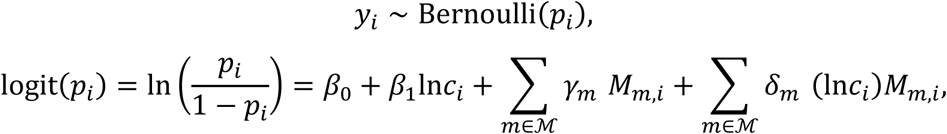

where *c*_*i*_ is the number of sign-consistent simple paths from the perturbed gene to the phenotype node for test *i*; *M*_*m,i*_ = 1 if test *i* used method *m* and 0 otherwise; and ℳ = {Algebraic, ODE, RWR} indexes the non-reference methods, with the signed shortest-path baseline absorbed into the intercept *β*_0_ and the baseline slope *β*_1_. The coefficient *β*_1_ gives the change in log-odds of a correct prediction per natural-log-unit increase in path count for the baseline, the γ_*m*_are method main effects, and the interaction terms *δ*_*m*_ therefore measure how each propagation method’s slope against path count differs from that of the signed shortest-path baseline.

### Software Environment

All agents are powered by an LLM, and complete agent traces and logs are released alongside the code. We implemented the FLASH-P agent team for Anthropic’s Claude Code CLI environment^32^, and used the Opus 4.7 model with a Anthropic Max account for all runs. Benchmarking alternative models was not our objective, and the architecture imposes no restriction on model choice, so it can equally be run with other frontier LLMs such as GPT or Gemini. We also evaluated locally-hosted open alternatives (Gemma4 31B, Qwen3 235B) and found that neither could sustain the full pipeline in a single zero-shot run, losing focus across the sequential agent calls. We therefore disabled end-to-end agentic execution for local models and tested each stage in isolation, with papers supplied as pre-downloaded PDFs to bypass unreliable local web fetching. Even under this favourable setup, the Builder, Refinement, and Reconciliation stages performed substantially below frontier-model output (Supplementary Note 1; Supplementary Fig. 1, 2). We did not test the largest open-weight models such as Llama 3.1 405B, DeepSeek-V3 671B, or Qwen3 480B, as the infrastructure required to host them locally is not widely accessible and was beyond our setup.

We designed FLASH-P with the emerging agentic AI ecosystem explicitly in mind, building it to accommodate plugins, external tools, and connections to other AI products, recognising that a scientific tool should not exist in isolation but should be interoperable with the growing range of AI-powered services available to researchers.

Runs were executed on a Dell Latitude 5550 laptop with an Intel Core Ultra 7 165H processor (1.40 GHz base, 16 cores) and 32 GB of RAM under Windows 11.

## Data Availability

All study data including network models, curated edges, perturbation datasets, Cytoscape files, and agent prompts are deposited in (https://github.com/CMits/FlashP). All resources are linked at https://flash-p.com.

## Code Availability

The FLASH-P framework software, code and analysis scripts are available on GitHub (https://github.com/CMits/FlashP). These results were generated with an earlier version of FLASH-P. Because agentic AI moves so quickly, we are continually evolving the agent, but this version was already performing well at the time it was used. The current version includes several further improvements, most notably greater token efficiency and more consistent network construction, with successive runs now recovering networks of largely the same core structure rather than producing substantially different topologies each time.

## Acknowledgements

This work was conducted by The Australian Research Council Centre of Excellence for Plant Success in Nature and Agriculture (CE200100015) and funded by the Australian Government. The project was also supported by an ARC Georgina Sweet Laureate Fellowship FL180100139.

## Author Contributions

D.K, C.M, C.B, N.F contributed to the main concept of the work. C.M developed the FLASH-P agents and software. D.K and C.M tested the software and produced results. C.M and D.K wrote and edited the manuscript.

---

**Extended Data Fig. 1.**
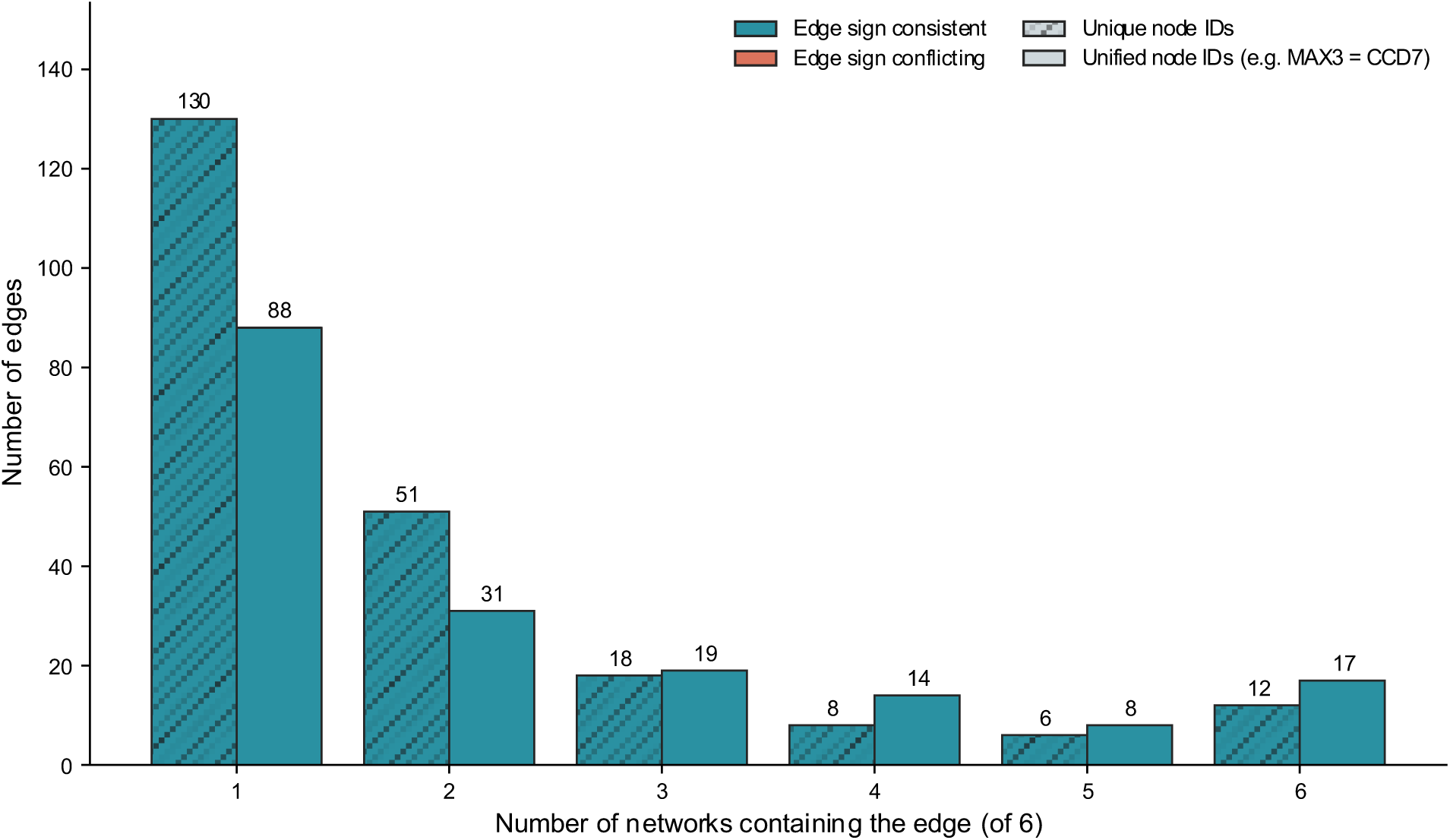
Edge recurrence across six independent FLASH-P runs for Arabidopsis shoot branching. Number of directed edges plotted against the number of networks (of six) in which each edge appears, under strict name matching (unique node IDs, hatched) and after alias resolution (unified node IDs, solid; e.g. MAX3 = CCD7). Bar colour denotes edge-sign agreement among networks sharing an edge (Blue, consistent; red, conflicting); no conflicting edges were found under either scheme. Most edges are run-specific, with 88 of 177 unified edges occurring in a single network, and unification raises overlap at the higher tiers without removing this predominance.

**Extended Data Fig. 2.**
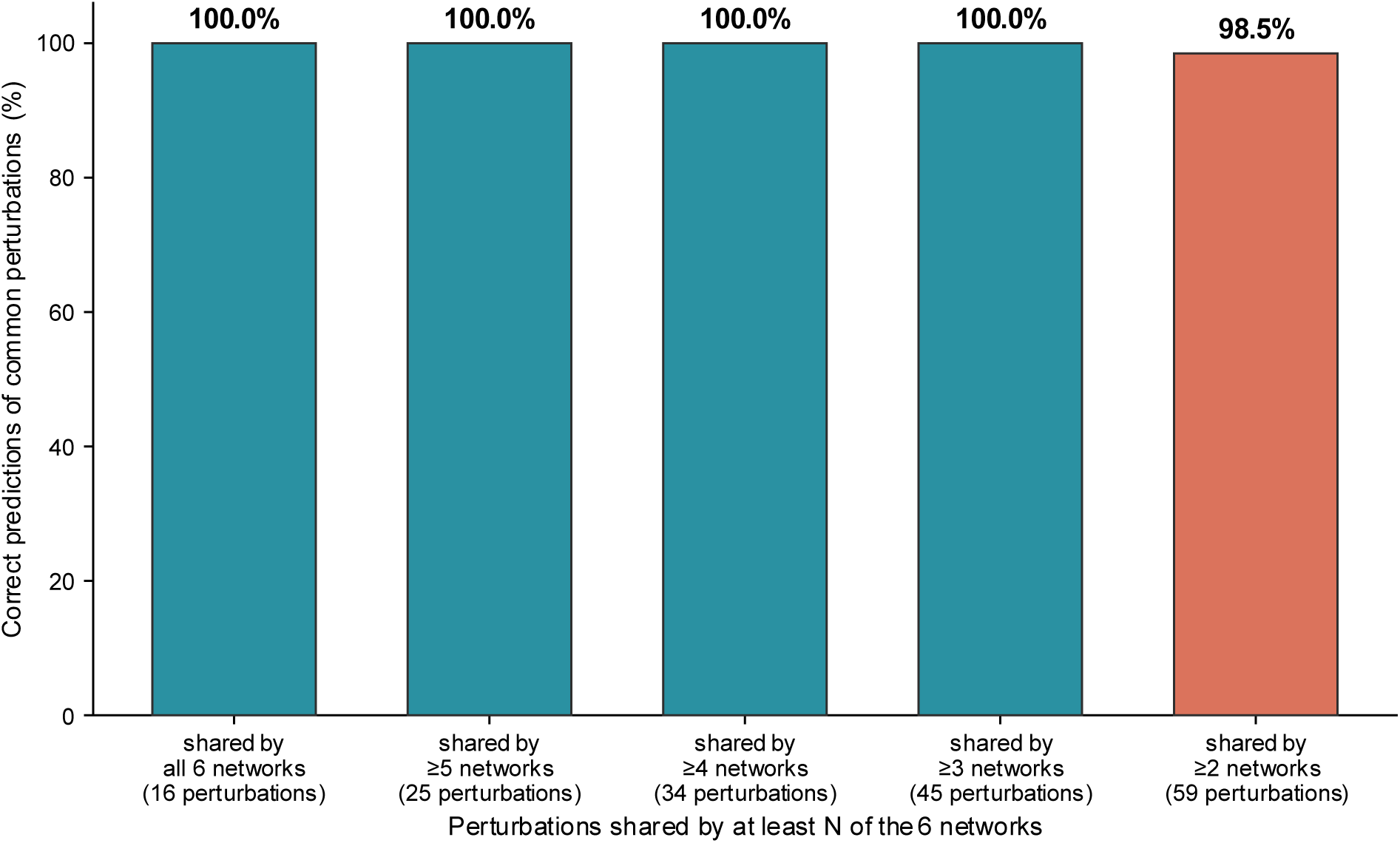
Predictive accuracy on perturbations shared across runs. Classification accuracy on perturbations common to at least N of the six networks, from all six (16 perturbations) to at least two (59 perturbations). Accuracy is the fraction of perturbation experiments for which the predicted direction of change at the shoot-branching node matches the experimentally reported outcome. Predictions are fully concordant for perturbations shared by three or more networks (100%) and remain high for those shared by at least two (98.5%).

**Extended Data Table 1.**
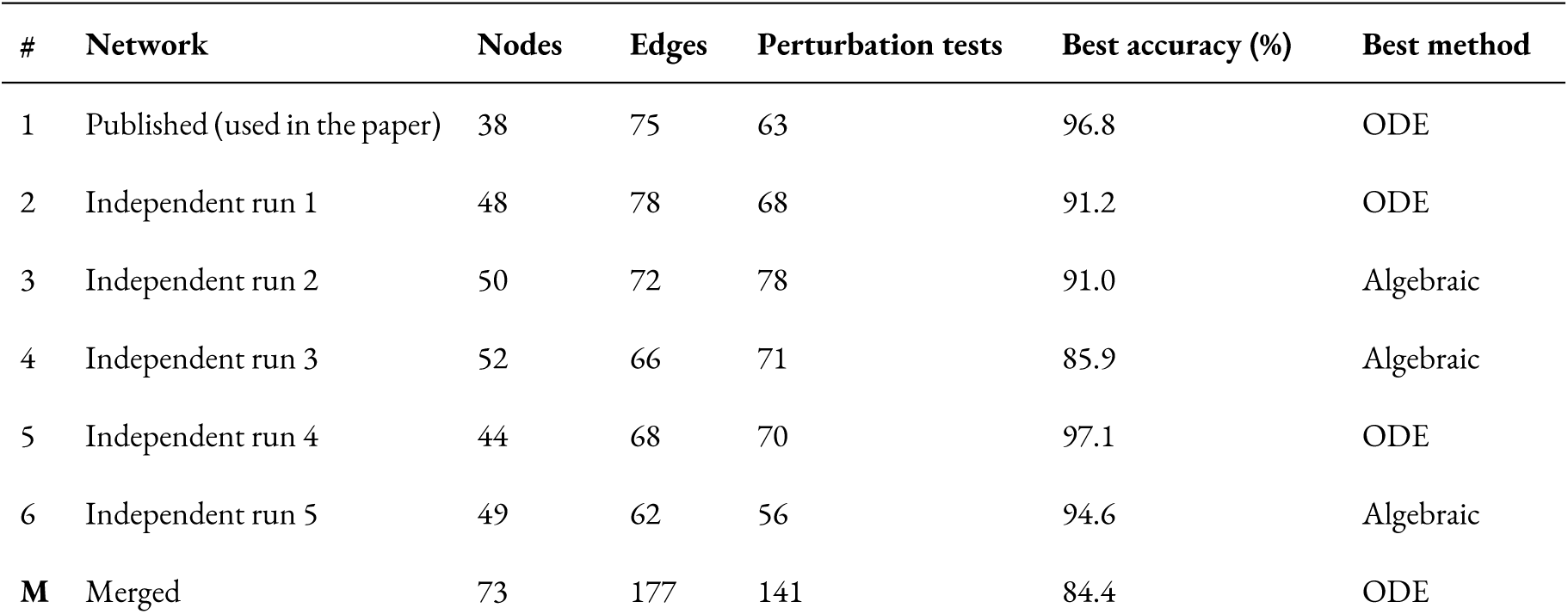
Reproducibility and consolidation of FLASH-P–constructed causal networks for Arabidopsis thaliana shoot branching, benchmarked against literature-reported perturbations. Each row reports one network built by FLASH-P for the same species–trait combination: the published network used in the paper (network 1) and five additional networks generated by independent, identical re-runs of the pipeline (networks 2–6), together with the merged ‘extended’ best-of-six network — the alias-deduplicated union of all six (final row, M). For each network the table lists the number of nodes and directed edges retained after construction, the number of perturbation tests reconciled from the literature (Tests), the highest classification accuracy obtained among the three network-propagation methods (Best Acc.), and the method that produced it (Best Method). The three methods are algebraic propagation, normalised Hill ODE, and random walk with restart (RWR). Accuracy is the fraction of perturbation experiments for which the predicted direction of phenotypic change at the trait node (shoot branching) matches the experimentally reported outcome. The five runs use an identical pipeline version, species and trait, differing only through the generative model’s stochasticity; the merged network is evaluated on the pooled union of all distinct perturbations extracted across the six networks.

| # | Network | Nodes | Edges | Perturbation tests | Best accuracy (%) | Best method |
| --- | --- | --- | --- | --- | --- | --- |
| 1 | Published (used in the paper) | 38 | 75 | 63 | 96.8 | ODE |
| 2 | Independent run 1 | 48 | 78 | 68 | 91.2 | ODE |
| 3 | Independent run 2 | 50 | 72 | 78 | 91.0 | Algebraic |
| 4 | Independent run 3 | 52 | 66 | 71 | 85.9 | Algebraic |
| 5 | Independent run 4 | 44 | 68 | 70 | 97.1 | ODE |
| 6 | Independent run 5 | 49 | 62 | 56 | 94.6 | Algebraic |
| M | Merged | 73 | 177 | 141 | 84.4 | ODE |

**Extended Data Table 2.** Node-identity alias map used to align the networks for comparison. Each row lists a unified (canonical) node and the original names that were treated as the same entity across the six networks and the human-curated model, with the type of relationship: Synonym (the same molecule under different names or abbreviations, e.g. MAX3 = CCD7); Paralog group (functionally redundant paralogs and their spelling variants merged into one node, e.g. SMXL6/7/8); Complex / family (a complex mapped to its functional subunit, or a redundant receptor/regulator family grouped, e.g. PIN3/4/7); and Compartmental (one entity modelled in several compartments integrated into a single node, e.g. shoot and bud auxin). Only nodes that required unification are shown; every other node appeared under a single name. To ensure the map does not manufacture agreement, all comparisons are also reported under exact (strict) name matching, which yields the same 100% edge-sign agreement.

| Unified node ID | Original names unified | Type |
| --- | --- | --- |
| CCD7 | CCD7, MAX3 | Synonym |
| CCD8 | CCD8, MAX4 | Synonym |
| CYTOKININ | CK, Cytokinin | Synonym |
| DECAPITATION | Decap_signal, Decapitation | Synonym |
| GA | GA, Gibberellin | Synonym |
| HIGH_R_FR | High_R_FR, Light | Synonym |
| LOW_R_FR | Low_R_FR, Shade | Synonym |
| STRIGOLACTONE | SL, Strigolactone | Synonym |
| SUCROSE | SUC, Sucrose | Synonym |
| BRC1 | BRC1, BRC1_2 | Paralog group |
| DRN_DRNL | DRN, DRNL, DRN_DRNL | Paralog group |
| FHY3_FAR1 | FAR1, FHY3, FHY3_FAR1 | Paralog group |
| HB21_40_53 | HB21, HB214053, HB21_40_53 | Paralog group |
| PIF4_5 | PIF4, PIF45, PIF4_5, PIFs | Paralog group |
| SMXL678 | SMXL678, SMXL6_7_8 | Paralog group |
| SPL9_15 | SPL15, SPL9, SPL915, SPL9_15 | Paralog group |
| ARR_A | ARR5, ARR_A | Complex / family |
| ARR_B | ARR1, ARR_B | Complex / family |
| CKX | CKX, CKX9 | Complex / family |
| IPT | IPT, IPT3, IPT7 | Complex / family |
| PIN347 | PIN3, PIN347 | Complex / family |
| YUC1 | YUC, YUC1 | Complex / family |
| AUXIN | Aux_Bud, Aux_Shoot, Auxin | Compartmental |

