## Supplementary Information for "FLASH-P: Turning decades of biology into accurate causal networks with AI agents"

**Supplementary Table 1. Network shallowness at the phenotype boundary in FLASH-P and KG-Cleaned *Arabidopsis* networks.** For each of the six individual phenotype networks (flowering time, hypocotyl length, lateral root density, plant height, seed size and shoot branching) and for the merged six-trait network, values report the percentage of directed edges whose target is a phenotype output node. Low percentages indicate deep networks in which most edges form intermediate regulatory cascades upstream of the phenotype output, and high percentages indicate shallow, star-like networks in which regulators connect to a phenotype node in a single step. FLASH-P networks are uniformly deep at the trait boundary (5.2 to 20.6% across the six individual networks and 10.9% in the merged network), whereas KG-Cleaned networks are dominated by direct phenotype edges (73.2 to 100% across the six individual networks and 92.5% in the merged network).

| network | phenotype | Edges directly targeting a phenotype node (%) |
|---|---|---|
| FLASH-P | Flowering_Time | 5.7 |
| FLASH-P | Hypocotyl_Length | 8.7 |
| FLASH-P | Lateral_Root_Density | 20.6 |
| FLASH-P | Plant_Height | 11.2 |
| FLASH-P | Seed_Size | 5.2 |
| FLASH-P | Shoot_Branching | 10.7 |
| KG-Cleaned | Flowering_Time | 100 |
| KG-Cleaned | Hypocotyl_Length | 73.2 |
| KG-Cleaned | Lateral_Root_Density | 99.7 |
| KG-Cleaned | Plant_Height | 100 |
| KG-Cleaned | Seed_Size | 100 |
| KG-Cleaned | Shoot_Branching | 100 |
| FLASH-P | merged_single_trait | 10.9 |
| KG-Cleaned | merged_pleiotropic | 92.5 |

### Supplementary Note 1. Citation provenance and hallucination audit

Every edge and perturbation in a FLASH-P network carries a machine generated literature citation (a DOI with a supporting sentence). We audited the provenance of these citations across all twelve networks in three stages. First we measured coverage. 98.8% of 2,464 edges and 100% of 1,521 perturbations carry a syntactically valid DOI rather than a placeholder. Second we tested resolvability by querying every unique DOI against the Crossref registry; 94.1% of the 644 unique DOIs resolve to a registered work, and the remainder are typographic or fabricated identifiers concentrated in a few networks. Per network coverage and resolvability are given in Supplementary Table 1.

Third we tested whether the cited paper actually supports the claim. From every network we drew a random sample (fixed seed) of four edges and four perturbations, giving 96 items. For each item an agent retrieved the cited paper, read the full text where it was openly available and otherwise fell back to the abstract, and judged whether the paper supports the stated claim, allowing for gene synonyms and family names. Reading the full text rather than matching gene tokens in the abstract removes the main weakness of an automated keyword check, namely that a paper can support a claim without naming the gene in its abstract. Per network results are given in Supplementary Table 2.

Across the 96 sampled items the cited paper was reachable in every case (42 via full text and 54 via abstract), and 89 of 96 (92.7%) were judged to support their claim. The 7 unsupported items were mostly cases where a correct fact had been attached to the wrong DOI.

**Supplementary Table 2.** Citation coverage and resolvability for the twelve networks. Edges and Perturbations are the counts curated per network, with the percentage carrying a syntactically valid DOI rather than a placeholder. Unique DOIs is the number of distinct identifiers cited in that network; Resolved (%) is the share that return a registered work from the Crossref registry and Not found is the count that do not. The all networks row counts each shared DOI once, so its Unique DOIs total is smaller than the column sum.

| Network | Edges | Edges with DOI (%) | Perturbations | Perturbations with DOI (%) | Unique DOIs | Resolved (%) | Not found |
|---|---|---|---|---|---|---|---|
| Flowering time (Arabidopsis) | 334 | 100 | 194 | 100 | 110 | 100 | 0 |
| Hypocotyl length (Arabidopsis) | 318 | 100 | 160 | 100 | 38 | 100 | 0 |
| Lateral root density (Arabidopsis) | 250 | 100 | 157 | 100 | 37 | 89 | 4 |
| Plant height (Arabidopsis) | 244 | 100 | 130 | 100 | 72 | 90 | 7 |
| Seed size (Arabidopsis) | 216 | 100 | 122 | 100 | 59 | 78 | 13 |
| Shoot branching (Arabidopsis) | 313 | 90 | 201 | 100 | 102 | 99 | 1 |
| Lycopene yield (E. coli) | 148 | 100 | 108 | 100 | 17 | 94 | 1 |
| Kernel row number (maize) | 146 | 100 | 77 | 100 | 43 | 100 | 0 |

| | | | | | | | |
|---|---|---|---|---|---|---|---|
| Lignin S/G ratio (poplar) | 141 | 100 | 34 | 100 | 42 | 98 | 1 |
| Tillering (rice) | 164 | 100 | 202 | 100 | 91 | 89 | 10 |
| Flowering time (sorghum) | 111 | 100 | 82 | 100 | 24 | 96 | 1 |
| Plant height (wheat) | 79 | 100 | 54 | 100 | 39 | 100 | 0 |
| **All networks** | **2464** | **98.8** | **1521** | **100** | **644** | **94.1** | **38** |

**Supplementary Table 3.** Agent verification of a random citation sample (four edges and four perturbations per network). Items checked is the sample size per network. Full text and Abstract only record how the cited paper was read. Supported counts items whose cited paper substantiates the claim; Not supported counts items reached but not substantiated (typically a real fact attached to the wrong DOI). Supported % is Supported divided by Items checked.

| Network | Items checked | Full text | Abstract only | Supported | Not supported | Supported % |
|---|---|---|---|---|---|---|
| Flowering time (Arabidopsis) | 8 | 4 | 4 | 8 | 0 | 100 |
| Hypocotyl length (Arabidopsis) | 8 | 4 | 4 | 8 | 0 | 100 |
| Lateral root density (Arabidopsis) | 8 | 7 | 1 | 8 | 0 | 100 |
| Plant height (Arabidopsis) | 8 | 3 | 5 | 7 | 1 | 88 |
| Seed size (Arabidopsis) | 8 | 0 | 8 | 7 | 1 | 88 |
| Shoot branching (Arabidopsis) | 8 | 3 | 5 | 8 | 0 | 100 |
| Lycopene yield (E. coli) | 8 | 8 | 0 | 8 | 0 | 100 |
| Kernel row number (maize) | 8 | 5 | 3 | 7 | 1 | 88 |
| Lignin S/G ratio (poplar) | 8 | 2 | 6 | 8 | 0 | 100 |
| Tillering (rice) | 8 | 2 | 6 | 6 | 2 | 75 |
| Flowering time (sorghum) | 8 | 1 | 7 | 7 | 1 | 88 |

| | | | | | | |
|---|---|---|---|---|---|---|
| Plant height (wheat) | 8 | 3 | 5 | 7 | 1 | 88 |
| **All networks** | **96** | **42** | **54** | **89** | **7** | **93** |

#### Supplementary Note 2. Local-model evaluation

Frontier large language models such as Claude Opus, GPT and Gemini are the default substrate for FLASH-P because the pipeline needs long context reasoning, faithful adherence to structured output schemas and reliable tool use across hundreds of sequential calls. Local open weight models are still attractive for groups that need to keep data on premises, avoid per call cost or remove dependence on third party infrastructure, so we asked whether a local model could sustain the full FLASH-P pipeline.

We ran all local experiments on a Mac Studio M4 Max with 128 GB of unified memory, using Ollama and contacting no proprietary cloud API. We tested two models. Gemma 4 31B was loaded with the Q4_K_M quantisation, 19 GB on disk and a 256K context window. Qwen3 235B-A22B was loaded with the Unsloth Q3_K_XL quantisation, 104 GB on disk and around 117 GB at inference, a mixture of experts architecture with 22B active parameters per token, a 256K context window, and native thinking mode and tool calling.

We first tried to run the pipeline autonomously, the same way FLASH-P runs with a frontier model. The local model received the FLASH-P agent specifications through the OpenCode tool calling harness and was asked to execute the steps end to end. This mode failed for both models. They could not use the tools efficiently at each step, particularly web search and web fetch during literature review, and they could not sustain the large running context, forgetting earlier outputs and losing track of what had already been done. We then checked web search and web fetch in isolation and the behaviour was equally poor, so we skipped that step and supplied the 68 paper PDF corpus used to build the unpublished shoot branching reference network directly as pre extracted text. This is the same corpus the frontier model used.

Even with the corpus supplied, neither model could orchestrate the rest of the pipeline autonomously. We therefore fell back to a prompt by prompt mode in which we invoked each step manually, copied the output of one step into the prompt of the next, and let a thin Python wrapper handle JSON parsing. This is the user assisted one by one execution referred to in the main text, and it is what distinguishes the local runs from the autonomous frontier model run. Because prompt by prompt mode removes essentially all of the orchestration overhead, the numbers below should be read as upper bounds on what these local models can deliver inside FLASH-P.

Single document edge extraction worked well in both models when each paper was supplied with a tight schema, and Gemma 4 processed all 68 PDFs in roughly four and a half hours, producing 390 edges and 175 perturbation tests with high schema compliance. Network construction is where the local models broke down. Gemma 4 produced a network with 12 nodes and 20 edges, substantially smaller than the reference network with 38 nodes and 75 edges, and it omitted the strigolactone receptor D14 together with the MAX1, MAX3 and MAX4 biosynthesis arm. Qwen3 235B did better in scope, with 31 nodes, 55 edges, and all canonical strigolactone components present. However, it introduced fourteen direct shortcut edges from upstream regulators to the Shoot_Branching node, bypassing the BRC1 hub. It also duplicated entities it had already represented, adding strigolactones, cytokinins, and other hormones as separate gene nodes, and inserting the MAX pathway as a standalone node despite already encoding its components. This reflects a fundamental failure in the builder's most important task which is maintaining awareness of what is already in the network and what is not.

We then ran a LLM driven perturbation reconciliation that added cross species ortholog mappings, signed RWR validation and a small refinement attempt. Gemma 4 reached 85.9% accuracy on its smaller test pool and Qwen3 235B reached 70.5% overall (93 of 132). Both models also failed at refinement. Across three iterations per model, neither produced a single iteration that improved accuracy by more than half a percentage point.

The accuracy of 85.9% from Gemma 4 is misleading because the network is very simple (Supplementary Figure 1), a central skeleton that a well prepared graduate student would draw from memory before reading any papers, with single hop cascades, no hubs, and no place for a breeder, gene editor or systems biologist to anchor downstream reasoning about pleiotropy, off target effects or candidate prioritisation. Qwen3 235B produces a substantially larger and visibly more complete network (Supplementary Figure 2) that does cover part of the strigolactone pathway, but expert review against the primary literature judged its biological quality below that of frontier model networks. For example, the BRC1 hub was bypassed by direct upstream to phenotype shortcuts, and the model was unable to revise its own structural choices when asked.

We conclude that frontier models remain required to run FLASH-P end to end at the quality reported in the main text, although larger open-source models were not tested due to infrastructure limitations.

#### Supplementary Note 3. Performance metrics

Each curated perturbation experiment is scored by comparing the predicted direction at the phenotype node against the experimentally reported outcome. Both the predicted and reported directions take one of three ordered values (decreased, unchanged, increased), indexed $k \in \{0,1,2\}$, and every test contributes one entry to a $3 \times 3$ confusion matrix $C$ in which $C_{ij}$ is the number of tests with reported class $i$ and predicted class $j$. We summarise this confusion matrix with four standard metrics, accuracy, Cohen's quadratic-weighted kappa, the multiclass Matthews correlation coefficient and per-class F1, alongside two bespoke metrics that combine prediction quality with network scope (FRS and DARS, defined below). Accuracy alone is class-imbalance sensitive and scale-blind, so the layered set is reported together on every row of the validator output.

Accuracy is the proportion of perturbation experiments for which the predicted class matches the reported class,

$$\text{Acc} = \frac{1}{T}\sum_{k} C_{kk} = \frac{\text{correct}}{T}, \qquad (1)$$

where $T = \sum_{ij} C_{ij}$ is the total number of tests. Accuracy is the easiest metric to read but also the easiest to inflate, because correctly predicting a single dominant class can produce a high value on an imbalanced test set.

Cohen's kappa corrects accuracy for chance agreement on the three-class confusion matrix. We use the standard quadratic-weighted definition for the ordered classes,

$$\kappa = 1 - \frac{\sum_{ij} w_{ij}\, O_{ij}}{\sum_{ij} w_{ij}\, E_{ij}}, \qquad w_{ij} = \frac{(i-j)^2}{(K-1)^2}, \qquad (2)$$

with $K = 3$ ordered classes, observed proportions $O_{ij} = C_{ij}/T$, and expected proportions $E_{ij} = (t_i/T)(p_j/T)$ under independent marginals, where $t_i = \sum_j C_{ij}$ is the number of tests reported as class $i$ and $p_j = \sum_i C_{ij}$ is the number of tests predicted as class $j$. Quadratic weighting penalises a decreased-versus-increased confusion four times more strongly than a decreased-versus-unchanged confusion, which is appropriate for an ordered output. $\kappa$ takes value 1 for perfect agreement, 0 for chance and is negative for systematic disagreement, and is the metric we use as a chance-corrected single-number summary on test sets in which the unchanged class is rare. We report 95% confidence intervals on $\kappa$ using the Fleiss–Cohen asymptotic standard error.

The Matthews correlation coefficient generalises the binary correlation between predictions and labels to the multiclass case in the standard sense,

$$\text{MCC} = \frac{c\,s - \sum_k t_k\, p_k}{\sqrt{(s^2 - \sum_k p_k^2)(s^2 - \sum_k t_k^2)}}, \qquad (3)$$

with $c = \sum_k C_{kk}$ the number of correctly classified tests, $s = T$ the total number of tests, and $t_k, p_k$ the row and column marginals defined above. MCC ranges from $-1$ for perfect disagreement through 0 for chance to $+1$ for perfect agreement, and is reported as a balanced summary that does not single out any individual class.

Per-class F1 scores are reported separately so that performance on the rare unchanged class is not hidden in the headline accuracy,

$$F1_k = \frac{2\,C_{kk}}{2\,C_{kk} + \sum_{j \neq k} C_{kj} + \sum_{i \neq k} C_{ik}}, \qquad (4)$$

which is the harmonic mean of precision and recall for class $k$, expressed directly through the confusion matrix. Bootstrap 95% confidence intervals on accuracy, $\kappa$, MCC and the per-class F1 are computed by resampling the $T$ per-test predictions 1,000 times with replacement.

To make networks of different size and test-set size directly comparable we additionally report the FLASH-P Rigor Score (FRS), a composite metric that combines chance-corrected prediction quality with scope. FRS is defined as

$$\mathrm{FRS} = \kappa \cdot \log_2\big(T \cdot (N + E)\big), \qquad (5)$$

where $\kappa$ is Cohen's kappa, $T$ is the number of validated perturbation tests and $(N + E)$ is the network size as the sum of the node and edge counts. The product $T \cdot (N + E)$ is the area of the validated-claim space, every test exercises a path through the network, so a larger network validated against a larger test set covers a larger region of biology, and the logarithm gives diminishing returns so that doubling the area adds a fixed quantum of credit. Multiplying by $\kappa$ ensures that this scope is only rewarded when predictions are better than chance; FRS is signed, so a network that is large but predicts at chance returns 0 and one that predicts worse than chance returns a negative value. FRS therefore reads as the chance-corrected number of bits of validated mechanistic claim that a network supports.

Because not all perturbation tests are equally hard, we also report a Difficulty-Adjusted Rigor Score (DARS) that replaces $T$ with a complexity-weighted effective sample size $T_{\mathrm{eff}}$,

$$\mathrm{DARS} = \kappa \cdot \log_2\big(T_{\mathrm{eff}} \cdot (N + E)\big), \qquad T_{\mathrm{eff}} = \sum_i c_i\,, \qquad (6)$$

where each test contributes a complexity score $c_i \in \{1,2,3\}$ defined as the number of mutations plus the number of treatments combined in that test (single perturbations score 1, double perturbations score 2, triple-and-higher perturbations score 3). DARS gives a network up to $\kappa \cdot \log_2 3 \approx 1.58$ extra bits of credit when its tests are uniformly hard, and an honest drop in $\kappa$ on hard tests cancels that bonus. Reporting FRS and DARS alongside accuracy and kappa allows scope, difficulty and quality to be read off a single row of the validator output without having to mentally trade off network size against test-set size or test difficulty.

We also report per-stratum kappa and accuracy on the easy / medium / hard partition (defined by the same complexity score) so that refinement can target the weakest stratum. Strata with fewer than five tests are flagged and the per-stratum kappa is suppressed to avoid reading noise as signal.

#### Supplementary Note 4. PlantConnectome export and cleaning

The KG-Cleaned baseline used in the main text is built from PlantConnectome[1], a knowledge graph that aggregates approximately five million functional relationships extracted by GPT-4o from over 71,000 plant biology publications. We exported one knowledge graph per Arabidopsis trait by querying the PlantConnectome REST API with phenotype-specific terms targeting the gene entity type. The terms used for each trait are listed in Supplementary Table 1.

**Supplementary Table 3. Seed search terms used to query PlantConnectome for each Arabidopsis trait.** All returned edges were aggregated into a single network per trait.

| Trait | Seed search terms |
|---|---|
| Shoot Branching | branching, axillary bud, tiller |
| Flowering Time | flowering, photoperiod, vernalisation |
| Hypocotyl Length | hypocotyl, elongation, photomorphogenesis |
| Plant Height | plant height, stem elongation, gibberellin |
| Lateral Root Density | lateral root, root branching, auxin transport |
| Seed Weight | seed size, seed weight, embryo development |

The raw PlantConnectome exports were not directly simulable because they contained contradictory edge signs, redundant entries, free-text node names with multiple gene identifiers and edges with unknown regulatory direction. We therefore applied a uniform cleaning script that retained only Arabidopsis-annotated edges, removed edges with unknown direction, normalised gene names onto the same nomenclature used by the FLASH-P networks, and mapped PlantConnectome relationship categories onto positive or negative regulatory signs. Positive (activation) categories included activation, induction, promotion, causation, positive regulation and explicitly positive results. Negative (inhibition) categories included repression, inhibition, decrease, negative regulation and suppression. Edges whose category was ambiguous (association, interaction or binding, all of which lack regulatory direction) were excluded.

The cleaned topology was then handed to exactly the same downstream pipeline that the FLASH-P networks use. Algebraic, normalised Hill ODE equations were auto-generated from the topology with the same script FLASH-P used, the same parameters and the same direction-classification thresholds. The head-to-head comparison in the main text therefore varies only the network's information architecture; the simulation and validation methodology is held the same as FLASH-P.

### Supplementary Figures

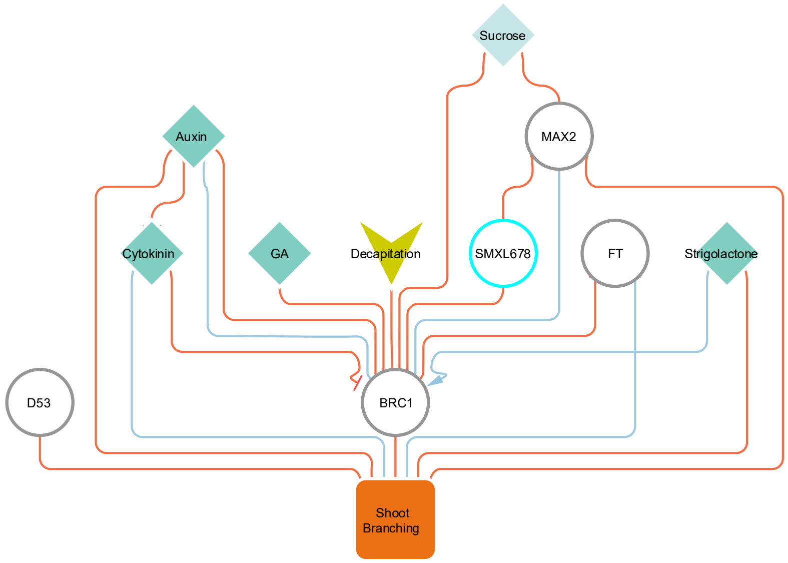


**Supplementary Figure 1:** Network produced for the Arabidopsis shoot branching corpus by Gemma 4 31B (Q4_K_M, served through Ollama on an Apple M4 Max workstation) under prompt-by-prompt user-assisted execution. The network has 12 nodes and 20 edges and lacks the strigolactone receptor D14 and the MAX1, MAX3 and MAX4 biosynthesis arm. Direction-call accuracy on the reduced reconciled test pool was 85.9 percent (55 of 64). The headline accuracy is misleading because the network covers only the most heavily reported core regulators and contains no peripheral hubs through which downstream applications such as breeding prioritisation or pleiotropy reasoning could be supported.

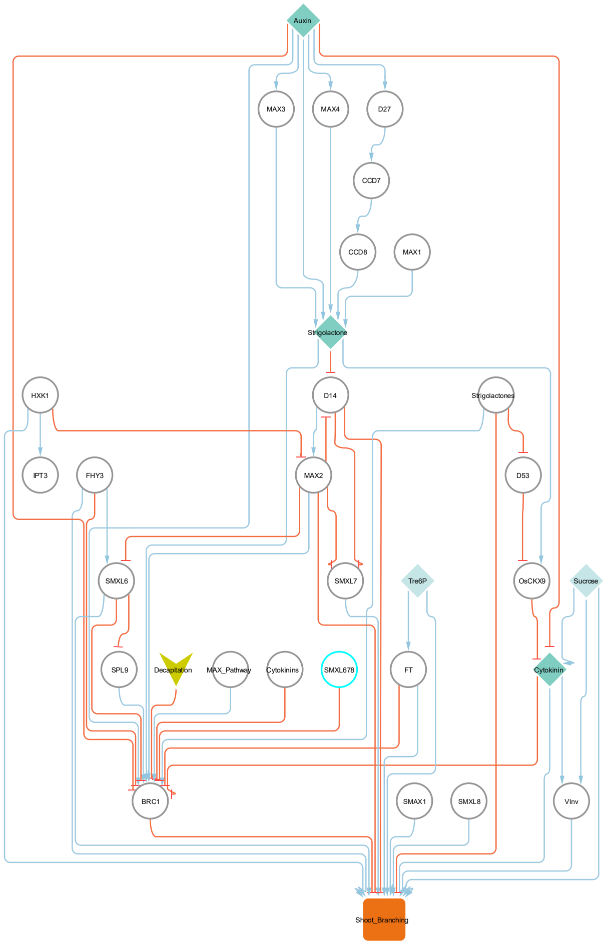


**Supplementary Figure 2:** Network produced for the same Arabidopsis shoot branching corpus by Qwen3 235B-A22B (Unsloth Q3_K_XL, served through Ollama) under the same prompt-by-prompt protocol. The network has 31 nodes and 55 edges and includes the canonical strigolactone components but introduces fourteen direct shortcut edges from upstream regulators to the Shoot_Branching node that bypass the BRC1 hub and prevent geometric-mean propagation through the cascade. Direction-call accuracy after LLM-driven perturbation reconciliation was 70.5 percent (93 of 132). Expert review of the network against the primary literature judged its biological quality below that of the frontier-model reference network, with edge-direction inconsistencies in places and a refinement step that did not improve accuracy.

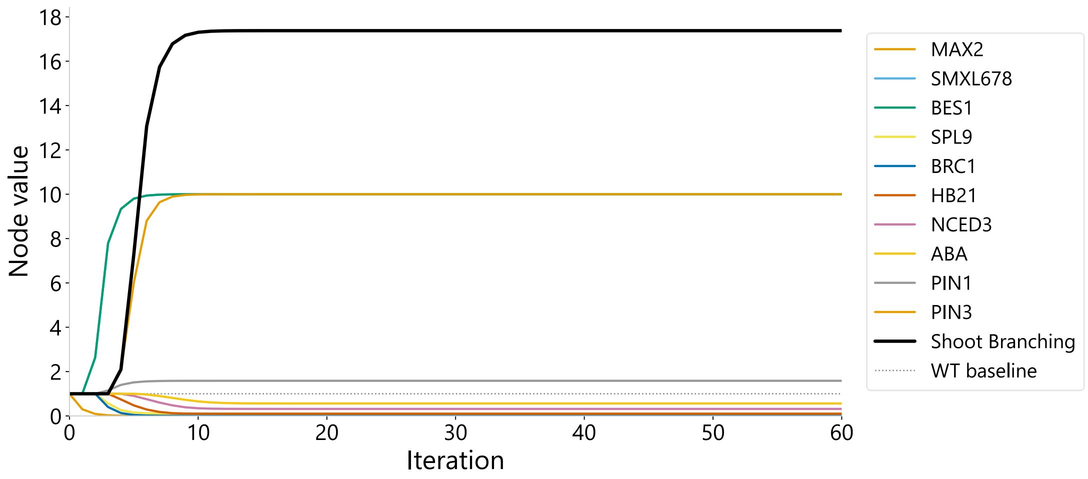


**Supplementary Figure 3:** Iteration-by-iteration trajectory of the eight nodes that move under the MAX2 knockout in the Arabidopsis shoot branching network, computed by the algebraic Jacobi solver with damping factor 0.7. The wild-type baseline is shown as a reference at value 1.0. Each trace converges to the steady-state value listed in Supplementary Table 3 by iteration 19; the trace is run to iteration 200 to expose the residual decay below the $10^{-4}$ convergence tolerance. This figure reproduces panel C of main-text Figure 2 and uses Supplementary Data 7 as its source data.

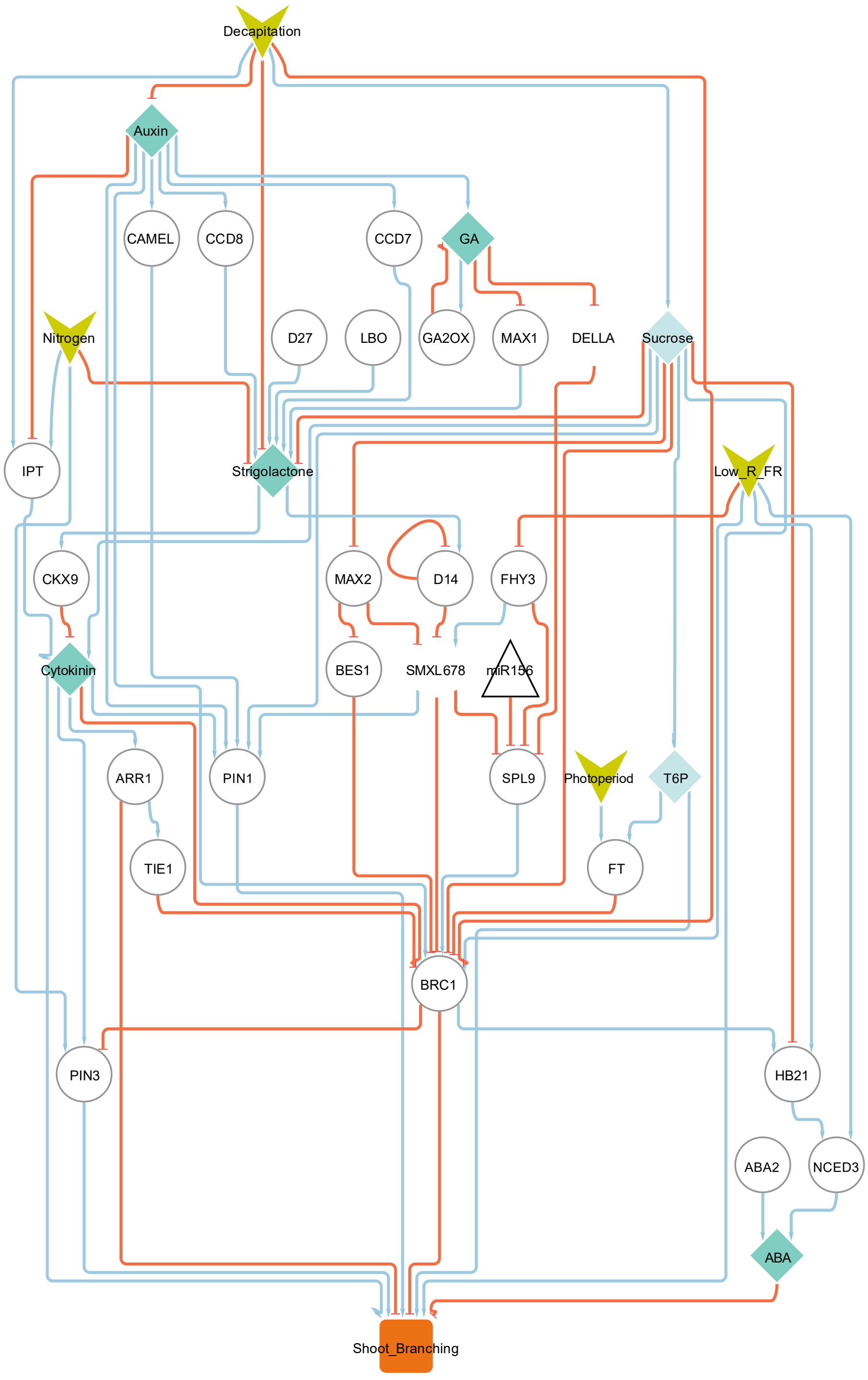


**Supplementary Figure 4:** Network created from Flash-P for Shoot Branching in Arabidopsis

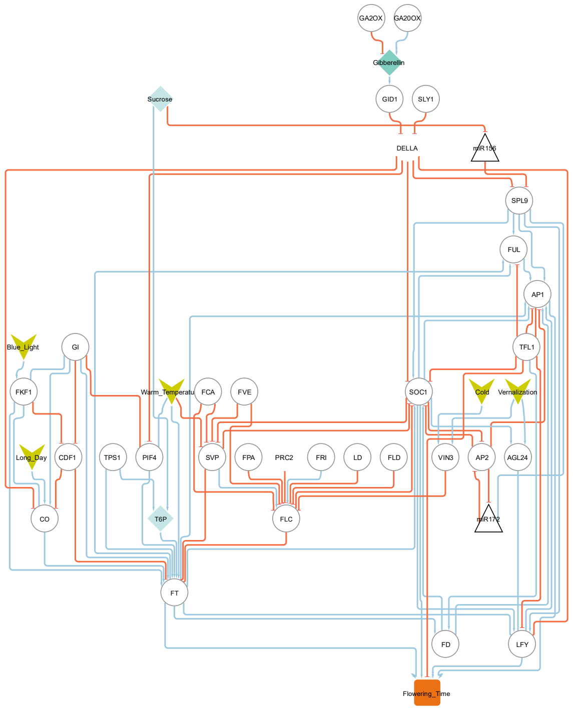


**Supplementary Figure 5:** Network created from Flash-P for Flowering Time in Arabidopsis

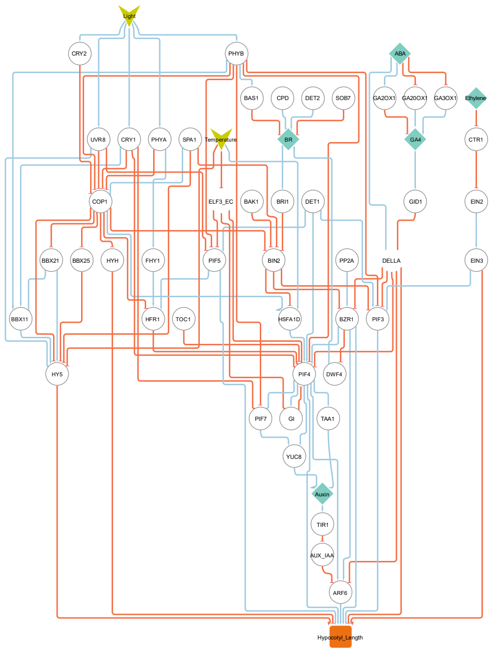


**Supplementary Figure 6:** Network created from Flash-P for Hypocotyl Length in Arabidopsis

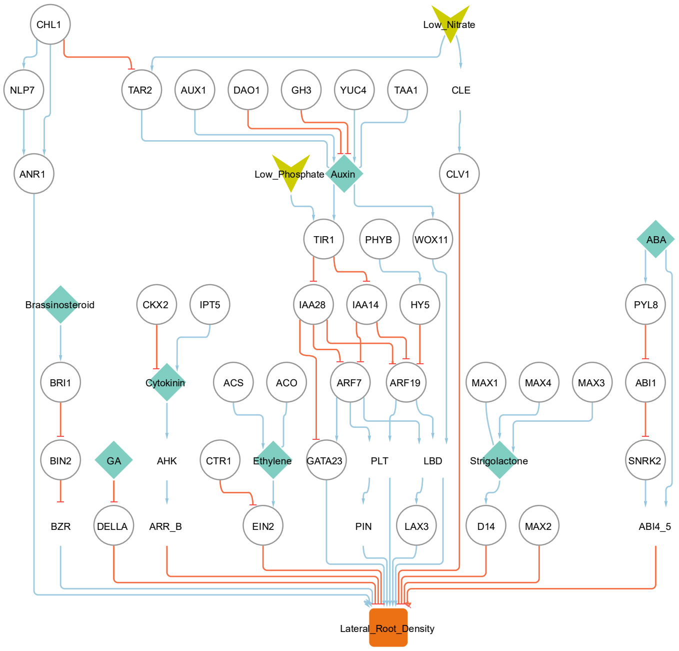


**Supplementary Figure 7:** Network created from Flash-P for Lateral Root in Arabidopsis

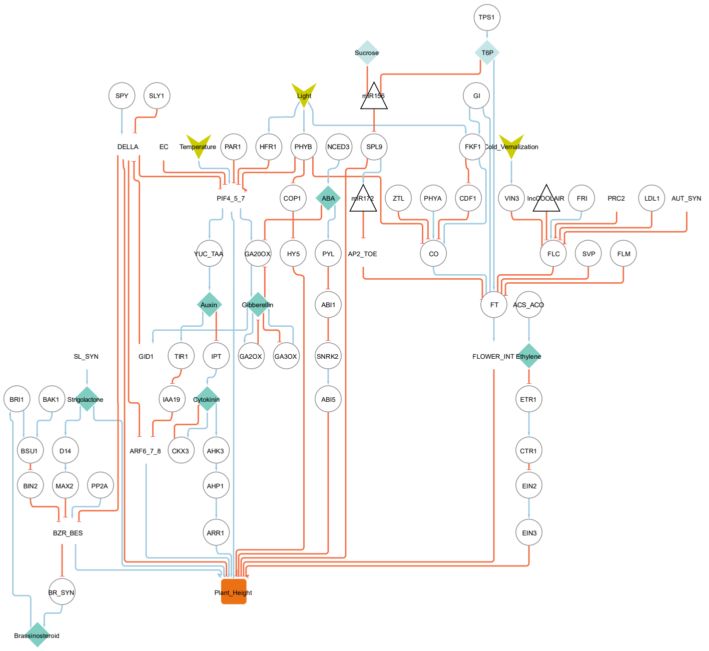


**Supplementary Figure 8:** Network created from Flash-P for Plant Height in Arabidopsis

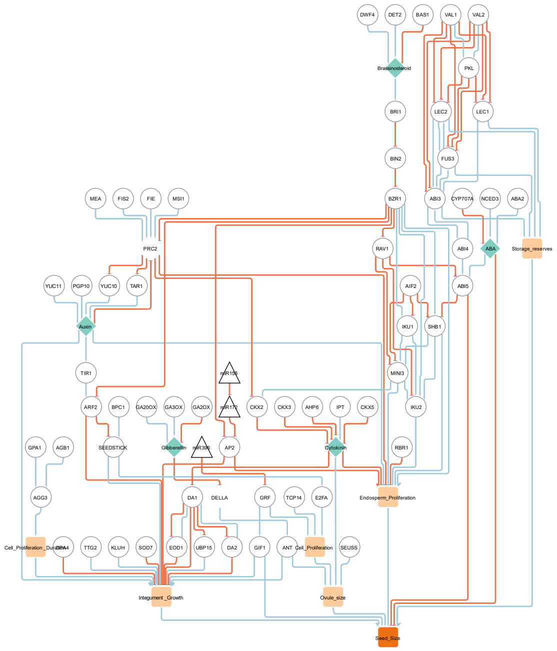


**Supplementary Figure 9:** Network created from Flash-P Seed Size in Arabidopsis

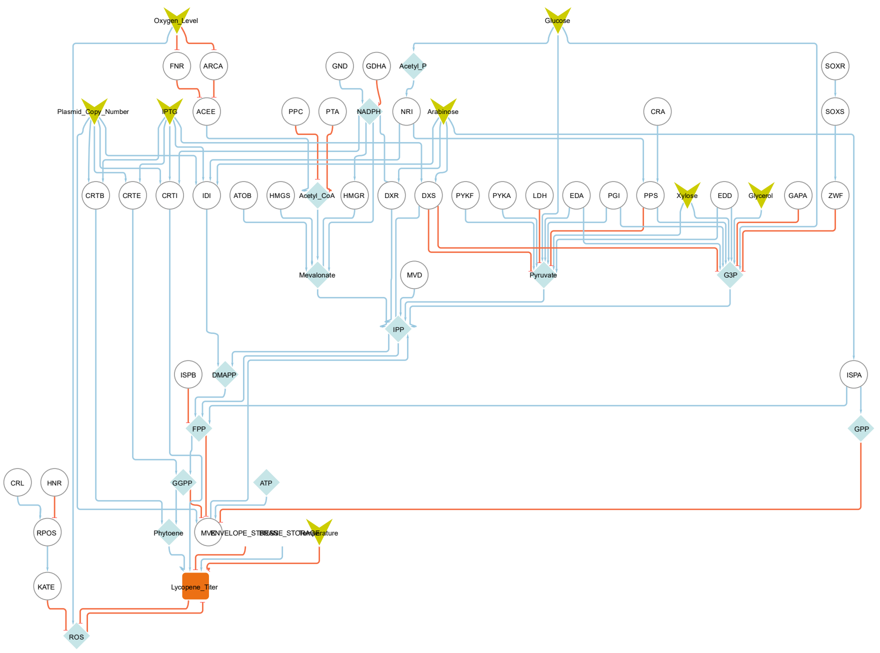


**Supplementary Figure 10:** Network created from Flash-P Lycopene production in E.coli.

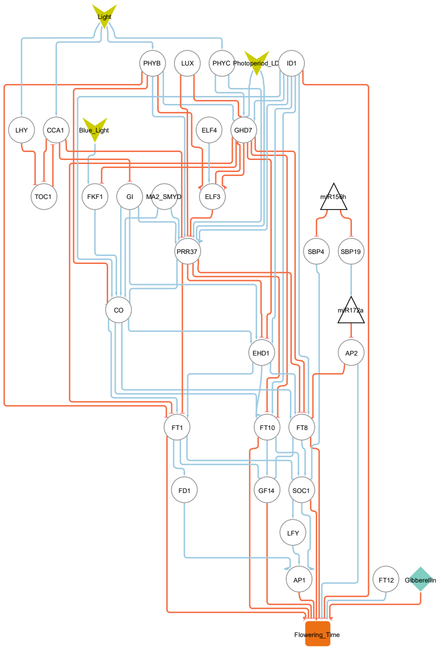


**Supplementary Figure 11:** Network created from Flash-P for Flowering Time in Sorghum

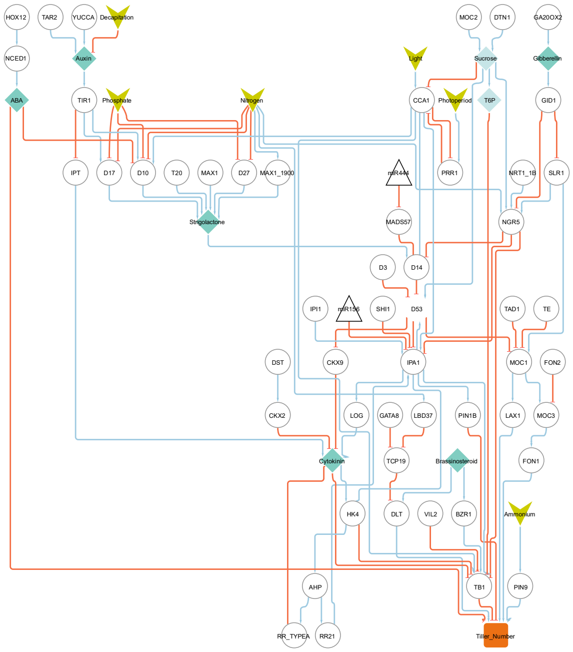


**Supplementary Figure 12:** Network created from Flash-P for Tillering in Rice

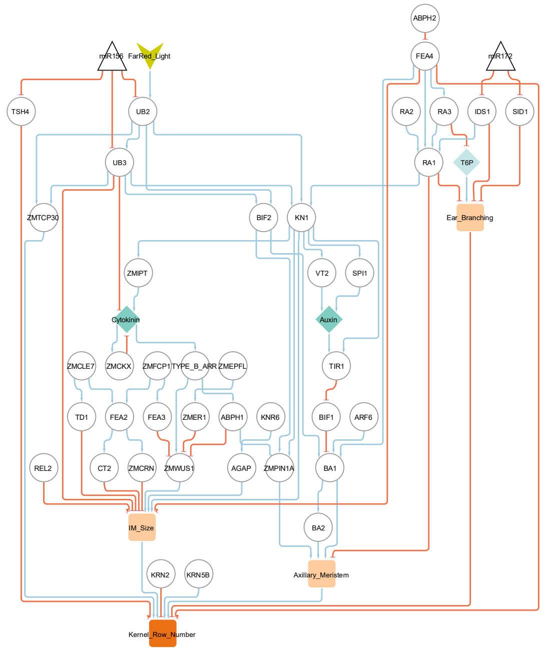


**Supplementary Figure 13:** Network created from Flash-P for Kernel row number in maze

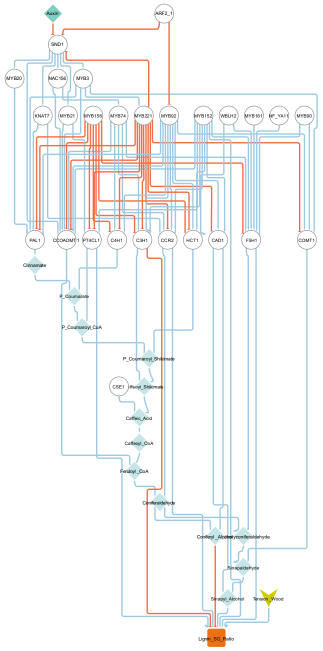


**Supplementary Figure 14:** Network created from Flash-P for S/G lignin ratio in Poplar

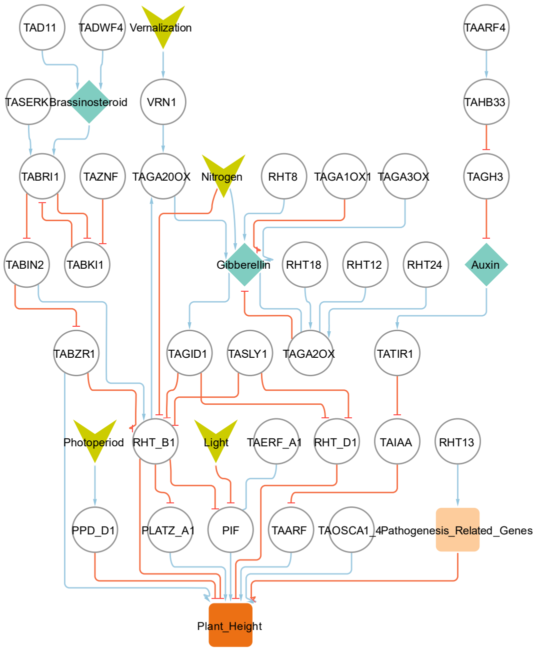


**Supplementary Figure 15:** Network created from Flash-P for Plant Height in Wheat

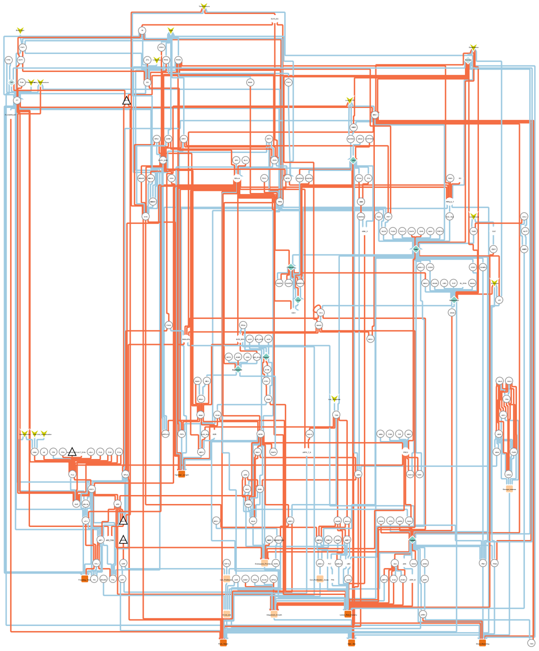

**Supplementary Figure 16:** Merged Network created from Flash-P from 6 traits of Arabidopsis created for this paper.
